# EpiSAFARI: Sensitive detection of valleys in epigenetic signals for enhancing annotations of functional elements

**DOI:** 10.1101/579847

**Authors:** Arif Harmanci, Akdes Serin Harmanci, Jyothishmathi Swaminathan, Vidya Gopalakrishnan

**Affiliations:** School of Biomedical Informatics, Center for Precision Health, University of Texas Health Science Center, Houston, TX, USA; School of Biomedical Informatics, Center for Systems Medicine, University of Texas Health Science Center, Houston, TX, USA; Department of Pediatrics, M.D. Anderson Cancer Center, Houston, TX, USA; Department of Molecular and Cellular Oncology, M.D. Anderson Cancer Center, Houston, TX, USA; Brain Tumor Center M.D. Anderson Cancer Center, Houston, TX, USA; Center for Cancer Epigenetics, University of Texas, M.D. Anderson Cancer Center, Houston, TX, USA; M.D. Anderson UTHealth Graduate School of Biomedical Sciences, Houston, TX, USA

## Abstract

The genomewide signal profiles from functional genomics experiments are dense information sources for annotating the regulatory elements. These profiles measure epigenetic activity at the nucleotide resolution and they exhibit distinct patterns along the genome. Most notable of these patterns are the valley patterns that are prevalently observed in many epigenetic assays such as ChIP-Seq and bisulfite sequencing. Valleys mark locations of cis-regulatory elements such as enhancers. Systematic identification of the valleys provides novel information for delineating the annotation of regulatory elements using epigenetic data. Nevertheless, the valleys are generally not reported by analysis pipelines. Here, we describe EpiSAFARI, a computational method for sensitive detection of valleys from diverse types of epigenetic profiles. EpiSAFARI employs a novel smoothing method for decreasing noise in signal profiles and accounts for technical factors such as sparse signals, mappability, and nucleotide content. In performance comparisons, EpiSAFARI performs favorably in terms of accuracy. The histone modification and DNA methylation valleys detected by EpiSAFARI exhibit high conservation, transcription factor binding, and they are enriched in nascent transcription. In addition, the large clusters of histone valleys are found to be enriched at the promoters of the developmentally associated genes.

## Introduction

Sequencing based functional genomics experiments, such as chromatin immunoprecipitation sequencing (ChIP-Seq), are being widely used for characterization of regulatory processes in the cells. Data generated from these experiments complement the DNA sequencing data for comprehensive understanding of genetic and epigenetic factors in disease(1) and in phenotypic variation(2). Recent years have brought about a substantial increase in the number of functional genomics assays(3–5). These assays generate a large trove of information that can be used for numerous purposes including PheWAS(6), HAWAS(7), and understanding the association between gene expression and epigenetic modifications(8, 9). In particular, these assays generate signal profiles that represent measurements at the signal nucleotide resolution and they are generally distributed in wiggle, bedGraph, and bigwig file formats(10). More specifically, the signal profile is a vector of values that represent a measurement for each nucleotide on the genome. In a ChIP-Seq experiment, the signal profile can be either the fold-change or the read coverage at each nucleotide. While the signal profiles may be affected by technical factors such as mappability, they contain much biological information that can provide biological insight for comprehensive discovery and annotation of functional elements in the genome.

Nevertheless, most of the current analysis pipelines focus mainly on the identification of broad regions that contain enrichment of signal. For example, peak calling identifies the enriched regions in ChIP-Seq data (11–14). While detection of the signal enrichments reveals the first layer of information about where enrichments manifest on the genome, the genomewide signal profiles from the functional genomics experiments contain much more information. In particular, there is much information that is encoded in the fluctuations of the signal profiles along the genome. Most notable of these fluctuations are the *troughs, valleys, or canyons* in the genomewide signal profiles. For example, the valleys within the read coverage signal profiles of histone modification (and transcription factor) ChIP-Seq data are indicative of open chromatin regions (i.e, nucleosome free regions) and of cis-regulatory elements such as enhancers.

In fact, valleys are commonly observed in the signal profiles generated from many epigenetic assays such as DNA methylation (bisulfite sequencing)(15, 16), open chromatin measurement (DNase sequencing)(17), and replication timing sequencing (for example, RepliSeq assay(18, 19)). In most of these cases, the valleys are the most important information bearing regions within the epigenetic signal profiles. Thus, the valleys can potentially enable researchers to anatomize and enhance the annotations of regulatory elements that can be identified from the epigenetic signal profiles. Consequently, the efficient and systematic identification of valleys can substantially increase the utility of functional genomics experiments. The information that are encoded in the valleys are currently left under-utilized because they are not reported explicitly by most of the analysis pipelines.

Several previous studies used valleys in the ChIP-Seq signal profiles(20, 21) for detection of regulatory elements such as enhancers. In most of these tools, technical factors such as mappability and nucleotide composition are not systematically considered in detection of valleys. As we discuss later, these technical factors may severely bias the location of the valleys. In addition, most of the tools are tailored for certain data types (e.g. ChIP-Seq data for histone modification of certain types) and do not generalize to other data types. Consequently, there is a dearth of generally applicable standalone methods that can be used to detect and analyze the valleys in genome-wide signal profiles. Here we present EpiSAFARI, which performs sensitive statistical detection of valleys from the fluctuations in the genome-wide signal profiles. The core algorithm is based on spline-based smoothing of the epigenetic signal profiles followed by valley detection using the smoothed signal profile. EpiSAFARI can analyze sparse signals such as DNA methylation signals which are non-zero only at cytosine residues on the genome. The kernel-based smoothing that is utilized by other methods is not applicable to the sparse signals because smoothing of the signal around missing values may potentially create artificial valleys. On the other hand, the spline-based smoothing of the signal utilized by EpiSAFARI does not suffer from this by ensuring that the fitting is performed only at non-zero values. An important technical variable for identification of valleys is that many valleys are caused by the low-mappability of the regions. For example, due to the fact that we cannot reliably map reads to genomic repeat regions, valleys may form around these repeat regions (14). These “non-biological” valleys may affect and bias the downstream analysis. Similarly, the nucleotide content of a region may impact the sequencing depth. For example, GC rich regions have been shown to be harder to sequence compared to the regions with uniform sequence composition(22). To take these factors into account, EpiSAFARI reports the nucleotide composition (count of each nucleotide and CpG dinucleotide frequency) and mappability statistics for all the reported valleys. The mappability and nucleotide content statistics are used to filter out any technical artefacts that may be introduced in the detection of valleys.

We first present the overview and main steps of the EpiSAFARI algorithm. We then present benchmarking of EpiSAFARI’s valley finding performance. We next demonstrate several applications of EpiSAFARI using DNA methylation and histone modification ChIP-Seq datasets. We analyze in detail the properties of histone modification and DNA methylation valleys with respect to conservation, transcription factor binding, DNA accessibility, and transcriptional activity.

## Materials and Methods

### EpiSAFARI Algorithm

EpiSAFARI algorithm is summarized in Figure 1. EpiSAFARI takes as input the mapped reads or the genome-wide coverage signals. The first step is smoothing of the signal using spline curves. EpiSAFARI generates a set of basis functions that are used to smooth the coverage signal (Supplementary Fig. 1). The spline-based smoothing is advantageous because they represent an efficient and effective method for noise removal(23). This step also decreases the number of false positive valleys that are created by the random fluctuations of genome-wide signal. EpiSAFARI divides the genome into windows of length *l_w_* and performs smoothing in each window. The smoothed signals are concatenated to form the genomewide smoothed signal. (See “Selection of Spline Parameters” Section, Supplementary Fig. 2). We observed that increasing *l_w_* too high may cause underfitting of the smoothed signal and may adversely affect valley detection. We evaluated the sensitivity of valley detection for selecting the spline smoothing parameters (knot numbers, knot location, and spline degree) and window lengths (Supplementary Fig. 3). We found that overly simple (small knot number and small spline degree) and overly complex smoothing parameters may cause underfitting and overfitting, respectively, and these may decrease accuracy.

**Figure 1:**
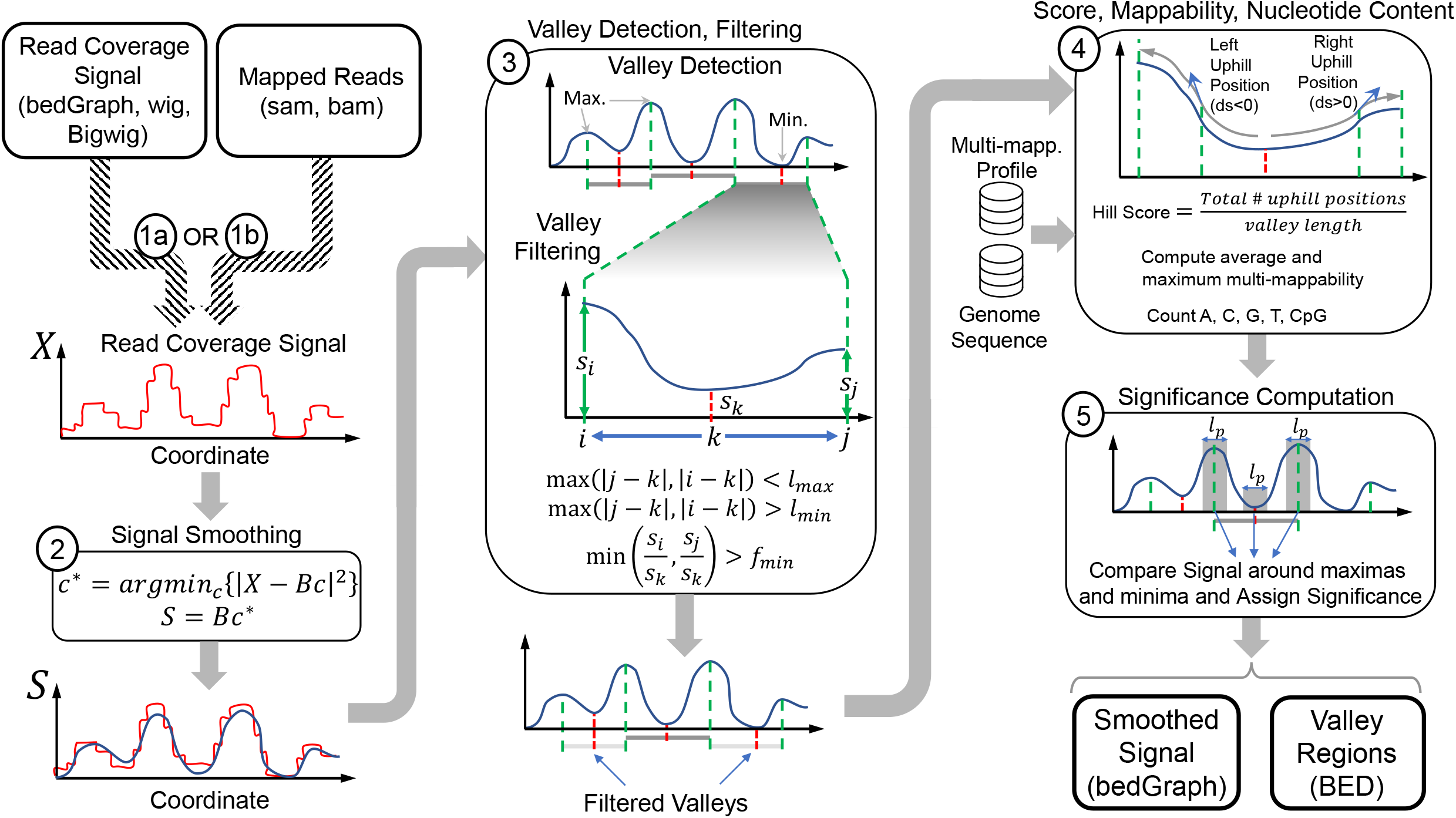
Illustration of the steps in EpiSAFARI algorithm. The input can be one of read coverage signal (bedGraph, wig, bigwig formatted) or mapped reads (sam formatted). The read coverage signal profile, *X*, is smoothed using spline-based fitting in the second step. Smoothing computes the minimum least square fit of *X* using the spline basis functions (*B*). The smoothed signal (*S*) is plotted with lightly colored original signal to illustrate the effect of smoothing. In third step, valleys are detected. Red and green dashed lines indicate the minima (dip) and maxima locations (summits), respectively. A valley is defined as a dip surrounded by two summits. The blow-up illustrates the summit positions (*i* and *j*) and the dip of a valley (*k*) and the signal level at these locations denoted by *s_i_, s_j_*, and *s_k_*. Next, the valleys are filtered with respect to distance between dip-to-summit distance (*l_min_, l_max_*) and the ratio between signal level at the summits to the dip (*f_min_*). The valleys that are removed are illustrated with grey shaded lines. In step 4, hill score, mappability, and nucleotide content are assigned. In this step, multi-mappability profile and genome sequence are used as input. In the final step, statistical significance for the valleys are assigned. For each valley, the signal enrichment is estimated using binomial test for comparing the signal within *l_p_* base pair vicinity of the summits compared to the signal around *l_p_* base vicinity of the dip. The output from EpiSAFARI are the smoothed signal profiles (bedGraph formatted) and the valleys (bed formatted)

After the signal is smoothed, EpiSAFARI identifies the local extrema, i.e. minima and maxima, locations in the signal. EpiSAFARI first computes the derivative of the smoothed signal and then identifies genomic positions where derivative changes sign from negative to positive as maxima (i.e., the summits) and positions where derivative changes sign from positive to negative as minima (i.e., the dips). These extrema are used to form the valleys. Each valley is defined as a region between two summit locations with a dip location that is between the summits (Fig.1). To decrease the search space of valleys, EpiSAFARI evaluates each dip and looks for two surrounding summits (one upstream, one downstream) within the *l_max_* base pair vicinity of the dip. In addition, the minimum distance between the dip and summit (*l_min_*) can be changed by the user. This parameter can be tuned if the user has certain expectation on the length distribution of valleys. In addition, changing *l_min_* and *l_max_* tends to increase the number of possible valleys quadratically because EpiSAFARI evaluates pairwise combinations all the summit pairs around each dip. EpiSAFARI also ensures that the fraction of the smoothed signal at each summit and the dip are higher than *f_min_*, the signal fraction cutoff. This way, EpiSAFARI ensures that the dip of the valley shows a certain depletion with respect to the summits.

After detection of the candidate valleys, EpiSAFARI computes several quality metrics and the statistical significance for each valley. These are used to filter out the valleys. First EpiSAFARI computes the multi-mappability and nucleotide content for each valley. The multi-mappability measures how well short reads can be mapped to a genomic location for a given read length(14). In general, high multi-mappability signal represents a lowly mappable region. These regions are especially important with regard to valleys because non-biological valleys (i.e., technical artefacts) can manifest around the regions with low-mappability. EpiSAFARI also computes the nucleotide content of each valley. For this, the counts of each nucleotide and the count of CpG di-nucleotides in the region are stored. The nucleotide counts also represent a technical factor similar to mappability because the regions with high (or low) GC content are known to be harder to sequence and non-biological valleys may be formed in these regions. In addition, the nucleotide counts are especially important when analyzing DNA methylation by Whole Genome Bisulfite Sequencing (WGBS). The user can use or exclude the valleys with high CpG and GC content in their analysis.

EpiSAFARI also computes a metric that we named “hill score” to evaluate the topological quality of the shape of valleys by assessing the signal profile on the two hills of the valley. A hill refers to the left or right side of the valley (Fig 1). The left (right) hill is the region between the left (right) summit and the dip location of the valley. A “good” valley has monotonically increasing hills while we are moving up on both of the hills. To measure this, EpiSAFARI starts moving from the dip to the left (right) summit and counts the fraction of locations where there is an up-hill trend. The left (right) hill score is the fraction of positions with up-hill signal to the length of the left (right) hill. The minimum of the hill score is used to filter out valleys.

EpiSAFARI finally assigns the statistical significance to the valleys. For each valley, the statistical significance, as computed by EpiSAFARI, reflects the difference between the signal enrichment at the summits and the depletion at the dip of the valley. EpiSAFARI first computes the total signal within *l_p_* base pair vicinity of the summits and the *l_p_* base pair vicinity of the dip location. EpiSAFARI then computes the significance of the depletion at the dip compared to the summits. We evaluated several background models for assigning the significance. The multi-mappability, nucleotide content, hill score and statistical significance are used to filter the valleys. We describe the details of the above step in detail.

### Spline-based Smoothing of Genomewide Profiles

The analysis steps of EpiSAFARI are illustrated in Fig. 1. The input to EpiSAFARI is the signal profile or the mapped reads. First step of EpiSAFARI is smoothing of the read coverage signal profile. To smooth the signal, EpiSAFARI divides genome into windows of length *l_w_* base pairs and applies spline-based smoothing to the raw signal profile in each window (Fig. 1, step 2). Although other smoothing approaches have been proposed(14, 24), basis spline curves are advantageous because they do not rely on a model, they are flexible, and they guarantee a continuous signal after fitting.

The spline curves are defined by a set of “knots” and the degree of the polynomial. The knots represent the positions where the polynomials meet such that the derivative of the curves are continuous up to the selected degree of the spline functions (Supplementary Figure 1). While knot positions affect the accuracy of spline-based smoothing, general knot selection is a complex and open problem(25). To study the effects of knot selection, we compared different knot selection procedures using different knot numbers and observed that uniform knot selection performs comparable in terms of accuracy to the random knot selection and derivative-based knot selection procedures (See “Selection of Spline Parameters”, Supplementary Figure 2a-c).

The degree of the splines represents the degree of the polynomials that make up the basis curves. By default, EpiSAFARI uses splines of degree 5 with 7 knots. We observed that increasing (or decreasing) the spline degree or knot numbers may decrease valley detection accuracy because they may cause underfitting or overfitting in the smoothing process (See “Selection of Spline Parameters”, Supplementary Figures 2a-c). The spline degree and knot number parameters tune the complexity of smoothing and they can be changed by the user. After basis function generation, a linear minimum square error fit of the signal to the spline basis functions is computed.

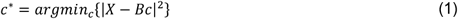

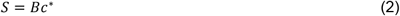

where *X* represents the vector that contains the original signal profile in the current window, *B* denotes the set of spline basis functions and *c*\* denotes the error minimizing weights. In order to decrease computational complexity, *X* is formed by using the signal levels at points of interest in each window. Specifically, these points are selected as the positions where signal changes value. For sparse signals (e.g. WGBS-based DNA methylation), the points of interest are chosen such that only locations with non-zero signal values are selected. Spline basis set represents a locally continuous set of curves that are used to smooth *X. S*, smoothed signal, is computed as the linear combination of the basis set weighted by *c*\*. It is important to note that the smoothing does not make any assumptions on the type of experimental assay. This enables EpiSAFARI to be applicable for analysis of the genome-wide signal profiles generated by any assay, for example microarray-based assays, i.e. ChIP-chip and DNA methylation arrays(26).

After smoothing, EpiSAFARI evaluates the maximum error among the points of interest. If the maximum error is higher than an anticipated error, EpiSAFARI increases the knot number and the spline degrees and fits the data with the updated parameters. This way, the spline parameters are automatically tuned to ensure that the curve-fit signal is close to the raw signal profile. As a result, even if the user sets out the fitting with a very simple model (i.e., low spline degree or small number of knots), EpiSAFARI will increase the complexity of smoothing parameters so that the observed error is smaller than maximum allowed error. After all the windows on a chromosome are processed, they are concatenated to form the final smoothed profile. In order to ensure that the transitions between consecutive windows are smooth, EpiSAFARI filters the concatenated signal profile using a short median filter. The length of the median filter can be changed by the user on the command line.

Some assays generate sparse signal profiles such that the signal levels are missing at arbitrary regions in the genome. An example of the sparse signals is DNA methylation by bi-sulfite sequencing(27) where the assays capture methylation levels for only the cytosine residues. The smoothing of these sparse signal profiles is challenging because we observe signal values mainly on cytosine (or CpG) residues. For the sparse signal profiles, EpiSAFARI utilizes only the non-zero signal values (i.e., only the positions that have signal value) for spline-fitting. For smoothing of DNA methylation signals, we set *l_w_* = 5000 base pairs and for smoothing the histone modification datasets, we set *l_w_* = 1000 base pairs (See Section “Selection of Spline Parameters”, Supplementary Figure 3a-c).

### Detection of Extrema in the Smoothed Profile

After smoothing the signal profile, next step is detection of the local extrema, i.e. local minima and maxima. EpiSAFARI first computes the derivative of the smoothed read coverage signal at each point. This is computed as the difference of signal between consecutive genomic positions as

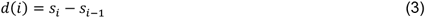

where *d*(*i*) represents the derivative of the signal at *i^th^* genomic position and *s_i_* denotes the smoothed signal value at *i^th^* genomic position. The minima are then identified as the genomic coordinates where derivative changes sign from negative to positive. Similarly, the maxima are identified as the locations where derivative changes from positive to negative. These are computed as

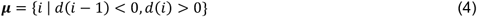

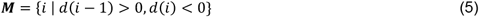

where ***μ*** and ***M*** represents the set of minima and maxima coordinates, respectively.

### Detection of Informative Valleys

After the local extrema are computed, EpiSAFARI detects the valleys. Each valley is a region defined by a local minimum (i.e. dip) and two nearby maxima (i.e., summits) located upstream and downstream the dip (Fig 1, step 3). The informative valleys are identified as valleys that have high difference between the signal value at the dip and at the summits. In addition, the summits are constrained to be within the local neighborhood of the dip:

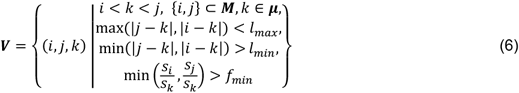

where ***V*** denotes the set of valleys, which are triplets of genomic positions (*i, j, k*). *i* and *j* denote the summit coordinates and *k* denotes the position of valley’s dip between *i* and *j* such that maximum distance from *i* and *j* to *k* are bounded by *l_max_* parameter and the minimum distance *i* and *j* to *k* are bounded by *l_max_* parameter. This ensures that the summits are not very far from (or near to) the dip. In addition, the ratio of the signal levels at *i* and *j* and *k* are bounded below by *f_min_*. This condition implies that there is difference between the signal levels at summits and signal at the dip. All the valleys that satisfy these constraints are reported. We set *f_min_* = 1.2 for sensitive detection of valleys in the experiments.

### Assignment of Hill Scores

For each valley, EpiSAFARI computes a quality score for the two “hills” on each valley. A hill is the genomic region between the dip and the (left or right) summits. A good hill shows a uniform increase between the dip and the summits (Supplementary Figure 4a). The measure this, EpiSAFARI computes the fraction of positions in the left and right hill where the signal is increasing (i.e., going up-hill) while moving away from the dip to the summit. This measures how uniformly signal is increasing within the hill. Hill score is computed as

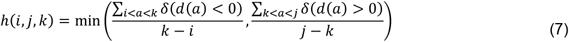

where *h*(*i, j, k*) denotes the hill score and *δ*(*d*(*a*) < 0) is an indicator function:

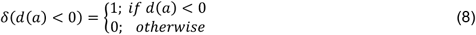

For a good valley, the hill score is close to 1.0, indicating that both hills to the left and right of the dip show a uniformly increasing signal while moving away from the dip. If there are any segment in the valley where there is down-hill trend, those regions decrease the valley score (Step 4 in Fig 1). Using a high hill score increases the topological quality of valleys but may adversely impact sensitivity of the valley detection. On the other hand, decreasing the hill score impacts the valley redundancy adversely (Supplementary Figure 4b). When the hill score threshold is low, smaller valleys start merging into larger valleys. This may complicate interpretation of the reported valleys since many valleys are overlapping with each other. We observed that the valleys with high qualities are separated in the distribution of the hill scores (Supplementary Figure 4e, f, g) at the very high end of the distribution. We therefore use high hill score cutoff to detect only the valleys with high topological quality.

The mappability of valleys is very important to distinguish valleys caused by low mappability versus the real valleys caused by biological signal fluctuation. For this, EpiSAFARI uses the precomputed multi-mappability signal(14) profile. For each valley, EpiSAFARI computes the average and the maximum of the multi-mappability signal. In general, high multi-mappability corresponds to a low mappable region and these regions can be filtered out in downstream analysis. EpiSAFARI reports the identified valleys in an extended BED file(28) (Supplementary Figure 6). For each valley, EpiSAFARI reports the location of dip of the valley and the summits, the signal levels at all the extrema, and the average and maximum of the multi-mappability signal on the valley. Another important factor for downstream analysis and filtering of valleys is the nucleotide content (i.e. GC content) within the valleys. EpiSAFARI computes and reports the per nucleotide counts and also CpG dinucleotide count within each valley. After valleys are processed, the user can view them on a genome visualization tool, such as IGV(29), concurrently with the signal profile.

### Assignment of Statistical Significance

The next step is assignment of statistical significance to the detected valleys (Fig. 1). With statistical significance, we would like to measure how significant the depletion of the signal at the dip is compared to the signal levels at the summits. Thus, the significance aims to measure the strength of the valley. The valleys with low p-value correspond to deep valleys. The valley’s significance is used to sort the valleys while performing enrichment analysis.

For a valley at (*i, j, k*), EpiSAFARI first computes the signal around the vicinity of the dip and the summits using

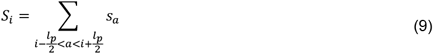

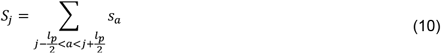

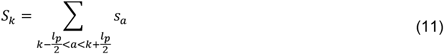

where *S_i_, S_j_, S_k_* denote the average signal in the *l_p_* base pair vicinity of the summits *i, j*, and the dip *k*. Next EpiSAFARI computes the binomial p-value of enrichment of signal around summits compared to the dip:

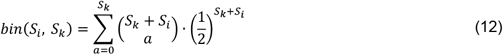

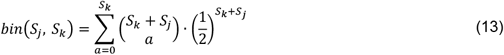

where 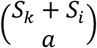 number of combinations for selecting *a* items within *S_k_* + *S_i_* items:

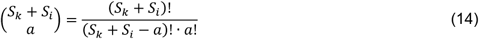

In order to assign the final p-value to the valley, we combine the p-values that are assigned to enrichment of the signal at the two summits. This process corresponds to combining the null models that are used to assign the two p-values for the observed summit-to-dip signal enrichment. We first use intersection of the null models as the joint null model (Supplementary Figure 7a). Assuming that the left and right hills are independent, this corresponds to the direct multiplication of the p-values:

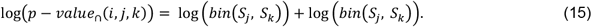

*p* − *value*_∩_ denotes the p-value computed by intersection-based combination of the p-values assigned to observed summit-to-dip signal enrichment. In addition, we use the union of the null models corresponding to null distribution of signal among summits and the dip so as to assign the p-value of the valley. As before, we assume that the p-values assigned to summits are independent from each other. Thus, the p-value estimated from the union of the null models is:

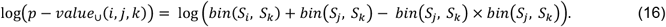

*p* − *value*_∪_ denotes the p-value computed by combining the individual p-values assigned to the observed summit-to-dip signal enrichments (Supplementary Figure 7a). In (15) and (16), we assumed that the p-values assigned to observed enrichment of the signal at the left and right summits are independent from each other. This assumption may not hold as we see a significant correlation of signals on left and right summits (Supplementary Figure 7c). As an alternative significance estimation method, we computed a multinomial distribution-based p-value assignment as this assigns a single p-value to the valley without the need for combining p-values. The multinomial p-value is computed as:

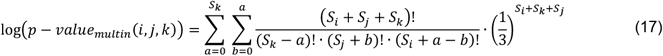

where the p-value is computed as the probability for different signal configurations at the summits and the dip such that the configurations are more extreme than what we observed. By more extreme, we mean the signal at one or both of the summits are higher than the observed signals. To compute this, we enumerate all the signal configurations such that the total signal at the dip, *S_k_*, is “distributed to the summits”. This way, all the possible signal configurations at the dip and summits is enumerated while the total signal is kept constant (Supplementary Fig. 7b). We then summated the probability of these signal configurations as they are computed under the multinomial distribution such that the probability of observing signals is distributed equally likely at the summits and the dip. The total probability corresponds to the multinomial p-value of the valley.

In general, we observed that union-based binomial p-value merging is much more conservative and shows lower sensitivity compared to the intersection-based p-value merging and multinomial based p-values (Supplementary Figure 7d, e). We therefore use intersection-based binomial p-value merging in the benchmarking. After the p-values are assigned, the false discovery rate at which each valley would be deemed significant is estimated using Benjamini-Hochberg procedure(30).

The valleys that EpiSAFARI detected may overlap with each other although we generally observed that the overlap between detected valleys tends to be very small. In order to get around this, EpiSAFARI can filter out the valleys whose dips are very close to each other. For this, EpiSAFARI goes through the valleys and detects the valleys whose dips are close to each other. For these valleys, EpiSAFARI selects the valley with the highest statistical significance (i.e., lowest p-value). This way the identified valleys represent sufficiently different dips in the signal (i.e., non-redundant valleys) and the most significant valley is detected around each dip.

### Selection of the Spline Parameters

The spline parameters determine the location of the valleys and may impact the accuracy. In addition, the hill scores and valley asymmetry may be affected by the spline smoothing parameters. Thus, we studied the impact of knot numbers, knot positioning, and spline degree on the accuracy of detecting valleys. To determine the effect of spline parameters on the valley accuracy, we used the H3K4me3 data for the K562 cell line from the ENCODE Project. To generate a ground truth set for the H3K4me3 valleys, we used the active genes promoters that overlap with any transcription factor binding peak as detected from ChIP-Seq datasets. The basic motivation for using these regions as ground truth is following: It is generally known that the promoters of active genes are enriched with H3K4me3 histone modification. If we put the additional requirement that these promoters overlap with transcription factor binding, these promoters most likely contain a valley inside them.

To generate the ground truth regions, we first downloaded the gene expression levels for K562 cell line from the ENCODE Project. We then identified the genes whose average replicate expression level is greater than 0.05. We finally extracted the promoters of these active genes and overlapped them with the ChIP-Seq transcription factor peak regions for K562 cell line from the ENCODE project. The intersecting promoters are used as the ground truth. We denote the genomic locations for the ground truth set of promoter regions by ***P***. It must be noted that these regions do not necessarily correspond to a complete set of the H3K4me3 valleys for K562 since this mark can also manifest on the enhancers in the intergenic domain. Thus, the valleys that EpiSAFARI detects will most likely contain many valleys that do not overlap with this ground truth. For this reason, we will evaluate the sensitivity of the valleys, i.e., the fraction of the set of ground truth regions that overlap with the detected valleys while evaluating the parameter selection.

After building the ground truth set, we next ran EpiSAFARI to detect the valleys in the H3K4me3 signal profile of K562 cell line with changing spline degree, knot number, and knot placement. To decrease the computation time, we focused only on the chromosome 1 for these analyses.

#### Knot Locations

To evaluate the effects of knot locations, we evaluated 3 knot placement strategies. First is derivative based knot selection. In this knot selection, we place the knots where the read depth signal shows fast changes along the genome. In this knot placement, EpiSAFARI places the knots at the locations for which the signal has the largest absolute signal derivative. We next implemented the random knot placement where the knots are randomly placed along the domain of the signal (Supplementary Fig. 1). We finally included the uniform knot placement where the knots are placed at equal intervals within the domain of the signal.

#### Knot Numbers

In order to include a wide range of knots distributed along the domain of the read depth signal, we used between 3 (minimum that we can use) and 15 knots. This way we evaluate both the densely and sparsely positioned knot selections. We denote the knot number with *k*.

#### Spline Degree

For each knot selection, we use spline degrees between 1 and 7. This way, we assess whether the increasing degree of the splines increase the sensitivity of the valley detection. We denote the spline degree with *ψ*.

We ran EpiSAFARI with the all the knot selection, knot number, and spline degree parameter combinations and computed the sensitivity of the identified valleys from each parameter combination. We next computed the sensitivity of the valleys as:

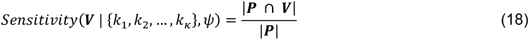

where ***V*** denotes the set of valleys (i.e., the genomic coordinates of the valleys) that are identified by EpiSAFARI with *κ* knots positions denoted by {*k*_1_, *k*_2_, …, *k_κ_*} and spline degree *ψ*. ***P*** denotes the set of active promoters (the genomic coordinates) that are bound by transcription factor peaks. |***P*** ∩ ***V***| denotes the number of active and TF bound promoters that overlap with the ***V*** and |***P***| denotes the number of promoters in ***P***.

We computed the sensitivity of the valleys detected using knot selection and spline degrees. (Supplementary Figure 2a, b, c). When the knot number and spline degree are both small, the sensitivity is smallest at around 0.2. As the number of knots or the spline degree increases, the sensitivities increase reaches around 0.8. This indicates that the overly simple smoothing is not powerful enough to detect the valleys. However, as we increase the complexity of smoothing, the sensitivity saturates at around 0.80 and starts decreasing as the smoothing is made more complex. This result highlights that increasing complexity of smoothing splines may decrease the sensitivity of valley detection. When different knot selection approaches are compared, the derivative based knot placement does not show improved performance over the uniform and random knot placement strategies. We also evaluated the number of valleys that are detected by EpiSAFARI using different parameter combinations (Supplementary Fig 2d, e, f). This is important because we want to also compare the number of valleys identified using different parameters. We observed that the number of valleys increases as the knot number and spline degree increases. The number of valleys (and sensitivity) decreases when we use parameter configurations with more than 7 knots and spline degree of 6 and higher.

In summary, we observed that extra complexity does not provide much improvement for our sensitivity analysis and in fact increasing complexity too much may cause overfitting of the data and may decrease the quality of selected valleys. Putting all these considerations together, we decided to use uniform knot selection with number of knots set to 7 and spline degree as 5. This selection is motivated to make balance between the accuracy, the number of valleys, and also the computation time that is required to run the algorithm (Supplementary Fig. 2g, h). The users can change the parameters to make EpiSAFARI run more conservatively or in a relaxed fashion.

It is worth noting that there are knot placement strategies other than the ones that we evaluated here(31). As we discussed before, the knot placement in spline smoothing is an open problem that is currently not solved in general cases. However, our results show that when we use a set of basis splines that are reasonably complex, i.e. not-very-low spline degree and knot numbers, the placement strategy does not impact the sensitivity considerably.

#### Window Length Selection

Window length parameter, *l_w_*, directly relates to smoothing as it determines the chunk of signals that will be smoothed at every step of smoothing. We computed the sensitivity of the detected valleys with changing *l_w_* parameter (Supplementary Fig 3). As *l_w_* < 1000, the sensitivity increases with increasing window length, after *l_w_* > 1000, the sensitivity starts decreasing. The main reason for this is possibly that the spline smoothing is underfit, i.e., the number of knots (and basis functions) is not large enough to reliably smooth the signal. From this observation, we suggest usage of *l_w_* = 1000 for punctate histone modifications. The selection of window length for sparse signals should be increased to increase the number of points of interest in each window so that the smoothing can be performed reliably. In addition, the expected valley lengths must be taken into consideration. For DNA methylation signals, we observed that for *l_w_* = 5000 is sufficient with the knot number of 7 and spline degrees of 5.

### Impact of Smoothing Parameters on the Hill Score

The hill score is computed for each valley separately using the smoothed signal profiles (Supplementary Figure 4a, b). Thus, the effect of the smoothing parameters on the computed hill scores is important. To compare the hill score estimates from different smoothing parameters, we computed the correlation between the hill scores assigned to valleys detected with different parameters. For this, we ran EpiSAFARI to identify the valleys in H3K4me3 data using the knot numbers, *κ*, between 4 and 15, and spline degrees, *ψ*, between 4 and 7. Given two sets of valleys computed by different knot numbers and spline degrees, we identified the valleys that share minima between these valley sets. Next, we computed the correlation between the left hill scores and the correlation between the right hill scores. This correlation computation is performed for all pairwise comparisons of parameters. The distribution of the left and right hill score correlations (Supplementary Fig. 4c, d) show that there is a substantial agreement between the assigned scores such that the correlations are mostly clustered above 0.40 with the most frequent correlations around 0.80.

It should be noted that the maximum allowed error in smoothing was set to a very large value while signal is smoothed in the above computations. This was performed to compare the impact of the parameters on the hill score without any parameter updates. When we decrease the maximum error in smoothing to the default values, we observed that the correlations between the assigned hill scores increases much. For example, the correlation of left and right hill scores for the most distant parameter sets (*ψ* = 4, *κ* = 4) and (*ψ* = 7, *κ* = 15) is 0.59 and 0.66, respectively. Whereas, without the parameter updates, the left and right hill score correlations between valleys detected from these parameter sets is 0.49 and 0.44, respectively. This indicates that the hill scores of valleys detected with the parameter updates will exhibit higher consistency.

#### Selection of hill score threshold with respect to sensitivity and valley redundancy

One of the important parameters is the hill score threshold that is used to filter out topologically low-quality valleys (Supplementary Fig. 4b). In principle, the higher hill scores correspond to valleys that have very good topologies such that hills are monotonically increasing as we move from the valley’s dip to the valley’s summits. Thus, setting the hill score threshold high enables selecting good valleys. The distribution of left and right hill scores (Supplementary Fig 4e, f, g) show that there are substantial number of valleys with hill scores very close to 1.

We next evaluated how the sensitivity of valleys changes with changing hill score threshold. We detected valleys using hill score parameters between 0.1 and 0.99. It can be seen that the sensitivity decreases as we increase the hill score. While the sensitivity of the valleys is decreasing with increasing hill score, another competing factor is the valley redundancy (Supplementary Figure 4k, l, m). The valley redundancy refers to how many valleys overlap with each other. The valley redundancy will increase with decreasing hill score because valleys may start engulfing other valleys when the hill score threshold is decreased. We computed the valley redundancy as:

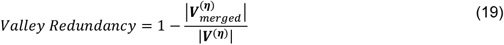

where *η* indicates the hill score, ***V^(η)^*** denotes the valleys detected using *η* and 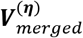 denotes the set of valleys generated by merging the valleys in ***V^(η)^*** where any two valleys with at least 1 base pair overlap are merged into one valley. As expected, the valley redundancy decreases with increasing *η* because valleys have distinctly uniform shapes. For *η* = 0.1, the redundancy is around 40% and decreases to around 3% for *η* = 0.99. This result indicates that hill score cutoff of 0.99 enables identification of distinct valleys at a cost of sensitivity. We have decided this is a fair tradeoff to generate high quality valleys and used *η =* 0.99.

### Impact of Smoothing Parameters on Valley Asymmetry

Similar to the hill scores, the smoothing parameters may impact the valley asymmetry, i.e., the imbalance between the left and right summits of the valleys. We first performed correlation of the valley asymmetry between every pairwise set of valleys within the sets of valleys detected using the knot numbers between 3 and 15, and spline degrees between 3 and 7 (Supplementary Fig. 5a). Most of the correlations are clustered around 0.9, which indicates a high consistency between the asymmetry of valleys detected by set of parameters. We also evaluated the fraction of valleys that “changed direction” when pairwise sets of valleys are compared. To detect valleys that changed direction, we compared pairs of valleys detected using different parameters, then we counted the number of valleys which have turned from a left-to-right valley to a right-to-left valley. By left-to-right (right-to-left) valley, we refer to the valleys whose left (right) summit has higher signal than the right (left) summit. The distribution of the fraction of valleys that changed direction (Supplementary Fig 5b) shows that directionality changing valley fraction is mostly clustered around less than 5%. These results indicate that the valley asymmetry is affected only slightly by the changing smoothing parameters.

### Valley Annotation

We extensively highlight the usage of transcription factor peaks for biological characterization of the valleys in the Results Section. EpiSAFARI can annotate valleys with respect to genes and transcription factor binding peaks. This step compares the valleys with an annotation file in GFF format and assigns the valley to the promoters, transcripts, and exons. We also created a GFF file from the transcription factor binding peak regions from ENCODE project(32). This GFF file contains the peaks of the transcription factors that are identified by 690 ChIP-Seq experiments performed on cell lines and uniformly processed by ENCODE project. EpiSAFARI can use these to annotate the valleys with respect to transcription factor binding. EpiSAFARI generates an extended BED file with the reported valleys and the spline-smoothed signal profiles in bedGraph format as output files. The extended BED file contains the valley positions, signal levels, multi-mappability signal, and annotations for all the valleys. The smoothed signal profiles can be used for visualizing the signal (Supplementary Fig. 6).

### Software Availability

EpiSAFARI source code is available upon request.

## Results

### Performance Benchmarking

We first focused on comparing the valleys detected by EpiSAFARI with the existing tools. While several studies have focused on analysis of valleys in different contexts, we found the PARE(21) is available as an algorithm that can be applied directly for comparison. It is important to note that PARE algorithm is designed as a tool for detection of the valleys with high specificity for the purpose of promoter and enhancer discovery. In the comparison, we used the H3K4me3 histone modification ChIP-Seq data for NA12878 individual from the ENCODE project(4). In general, H3K4me3 modification marks the promoters of the active genes. We have focused specifically on this modification because firstly it is a well-characterized mark and secondly PARE algorithm is tuned for analysis of this mark so that we are fair in comparison of the tools. We downloaded the two replicates that are available and pooled the reads from the replicates. PARE algorithm is run with default settings except that we extended the search space to 2,000 base pairs (-v option) and we relaxed the FDR cutoff to 0.1 (-t option). For EpiSAFARI, we set the FDR cutoff to 0.05, filtered out the valleys with hill scores lower than 0.99, and also filtered out the valleys for which the average multi-mappability is higher than 1.2. In general, we observed that EpiSAFARI identifies many more valleys compared to PARE. To make the comparison fair, we sorted the EpiSAFARI valleys with respect to increasing FDR (i.e., more significant first) and we sorted the PARE valleys with respect to decreasing score (i.e., higher score first) assigned by the algorithm. We then focused on the top 2,000 valleys. Since we do not have a set of valleys that can directly serve as ground truth, we used different hypotheses to evaluate whether the identified valleys are biologically meaningful and used these to assess performance of methods.

We first focused on comparison of transcription factor binding activity around the valleys. We hypothesized that the real valleys must be enriched in transcription factor binding. ENCODE project supplies a large number of ChIP-Seq experiments and uniformly processed peak calls for many transcription factors for NA12878 sample. We pooled the available peaks calls from the 90 ChIP-Seq experiments for NA12878 sample. We then evaluated the fraction of top valleys that overlap with a transcription factor peak. In order to correct for the valley lengths reported by the methods, we used the valleys that are reported by PARE as they are and we used only the 200 base pair vicinity of the valley dips (i.e., dip location +/− 100 base pairs) reported by EpiSAFARI. Figure 2a shows the fraction of top valleys that overlap with a transcription factor peak while the number of top peaks is increased (x-axis). We observed that more than 90% of the top EpiSAFARI valleys that we evaluated overlap with a peak. The fraction of overlap decreases slowly as we increase the number of top valleys. PARE valleys, in contrast, show a fairly low overlap with a transcription factor peak (starting at 50%) and the overlap fraction increases as the number of top valleys is increased. This result indicates that the valleys (and the scores) detected by EpiSAFARI represent a better representative set of transcription factor activity.

**Figure 2:**
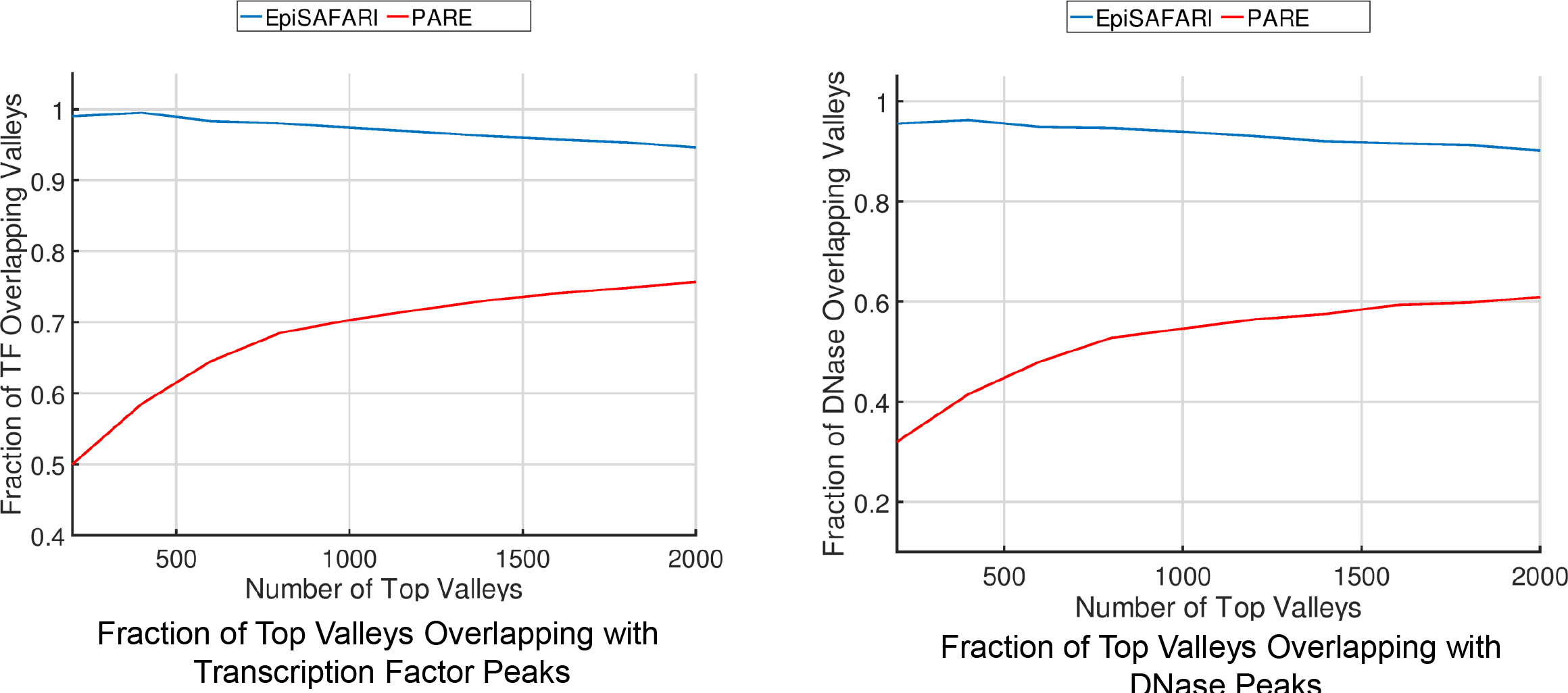
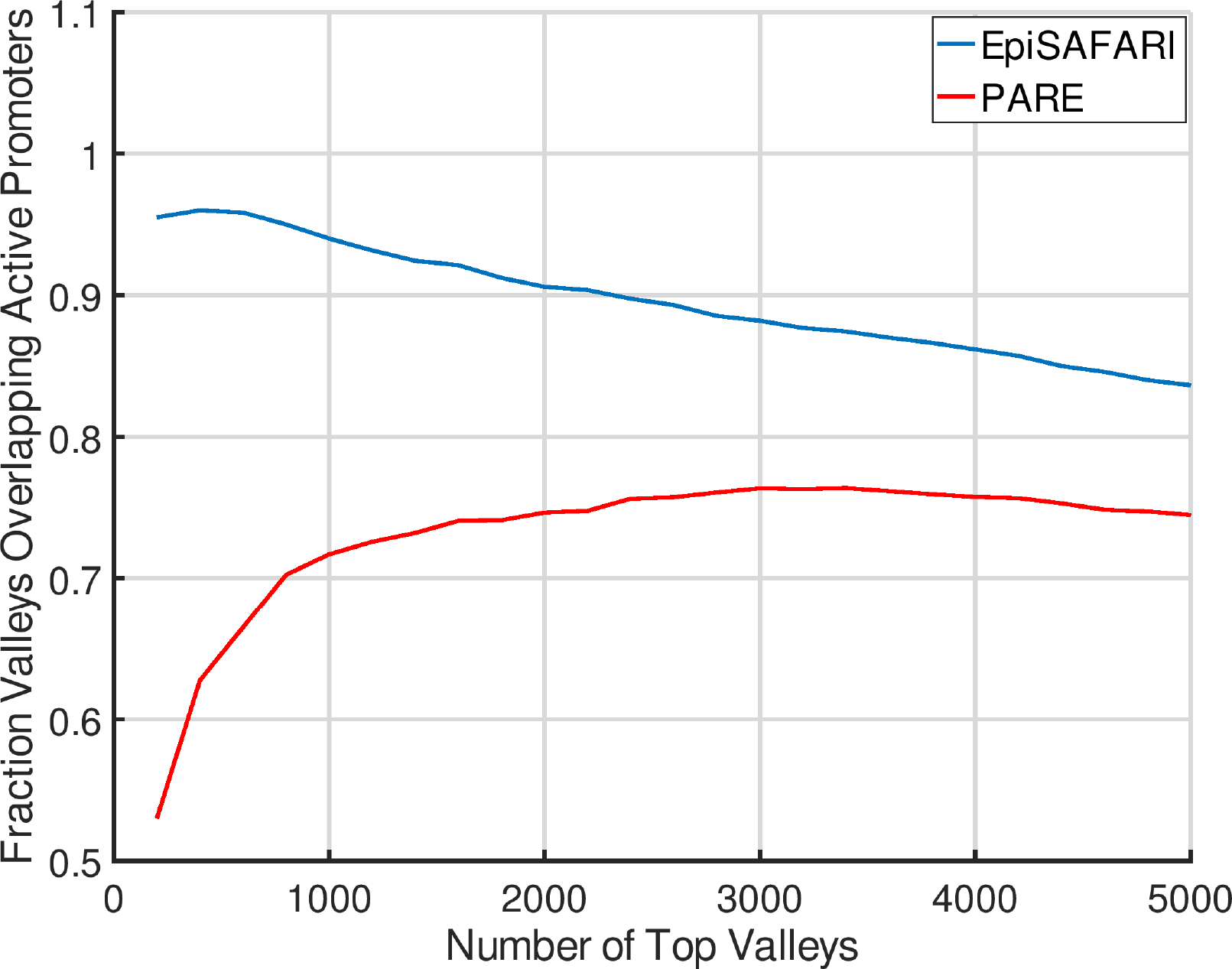
Comparison of the top H3K4me3 valleys in NA12878 sample as detected by EpiSAFARI (blue) and by PARE (red). a) The fraction of top valleys that overlap with a transcription factor peak. X-axis shows the number of top valleys and y-axis shows the fraction of valleys that overlap with a transcription factor peak. b) The fraction of top valleys that overlap with a DNase peak. c) The fraction of top valleys that overlap with an active promoter.

Another hypothesis about the valleys is that they are enriched in terms of open chromatin. To measure this, we downloaded the peaks of the DNase-1 hypersensitive site sequencing (DNase-Seq) data from the ENCODE project for NA12878 sample. These peaks represent the experimentally detected locations of genomic positions for accessible DNA. Similar to the previous comparison, we overlapped the top valleys detected by EpiSAFARI and PARE with the DNase peaks. Figure 2b shows the fraction of top valleys that overlap with a DNase peak. Similar to previous analysis, we used the 200 bp vicinity of the valley dip for EpiSAFARI valleys for this comparison. We observed that EpiSAFARI valleys show a much higher overlap fraction to the DNase peaks compared to PARE. In addition, PARE valleys also show an increasing overlap fraction with decreasing score while EpiSAFARI valleys show a slowly decreasing overlap fraction with decreasing significance. This result indicates that EpiSAFARI valleys are better representatives of the accessible DNA positions compared to the valleys detected by PARE.

We hypothesized that the top valleys of the H3K4me3 modification must be enriched in the active gene promoters. To identify the active gene promoters, we used the transcript expression quantifications from the ENCODE project for NA12878. We first identified the transcripts whose reported expression levels in terms of reads per kilobase per million mapped reads (RPKM) are higher than 0.05. We next extracted the 1000 base pair vicinity of the transcription start site of each transcript. These constitute the set of active promoters. We then overlapped the valleys with the active promoters and computed the fraction of top valleys that overlap with active promoters. Figure 2c shows the overlap fraction of top valleys with active promoters. EpiSAFARI shows a fairly high overlap (higher than 90%) at the top valleys and decreases as the number of top valleys decrease. For the top 2000 valleys, the overlap is always higher than 80%. The top valleys identified by PARE show 50% overlap with active promoters and the overlap increases as the number of top valleys is increased. This result shows that EpiSAFARI valleys capture the active promoter information better than PARE while detecting valleys.

### Delineation of Transcription Factors Binding within Histone Valleys

While we have evaluated the overlap of transcription factor binding to the valleys, it is also informative to study which specific transcription factors are bound at the valleys. For this, we overlapped the top 1000 valleys identified by EpiSAFARI with the peaks of each transcription factor separately and counted the number of overlaps for each transcription factor. Figure 3a shows the number of valleys that overlap with the peaks of transcription factors. As expected, the factors and complexes associated with active promoters are highly enriched. Among these Polymerase 2 (Pol2) peaks overlap with more than 95% of the valleys.

**Figure 3:**
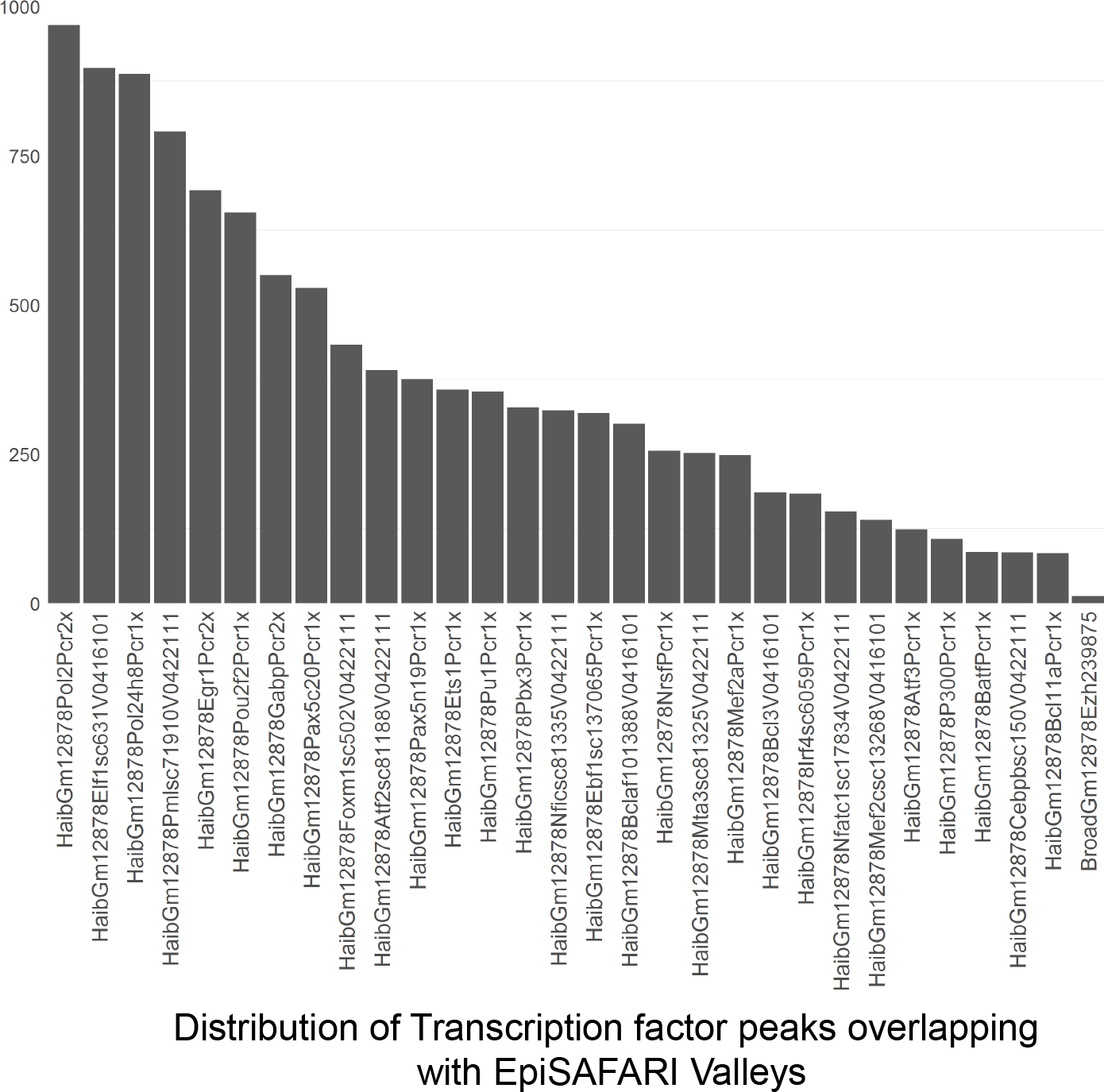
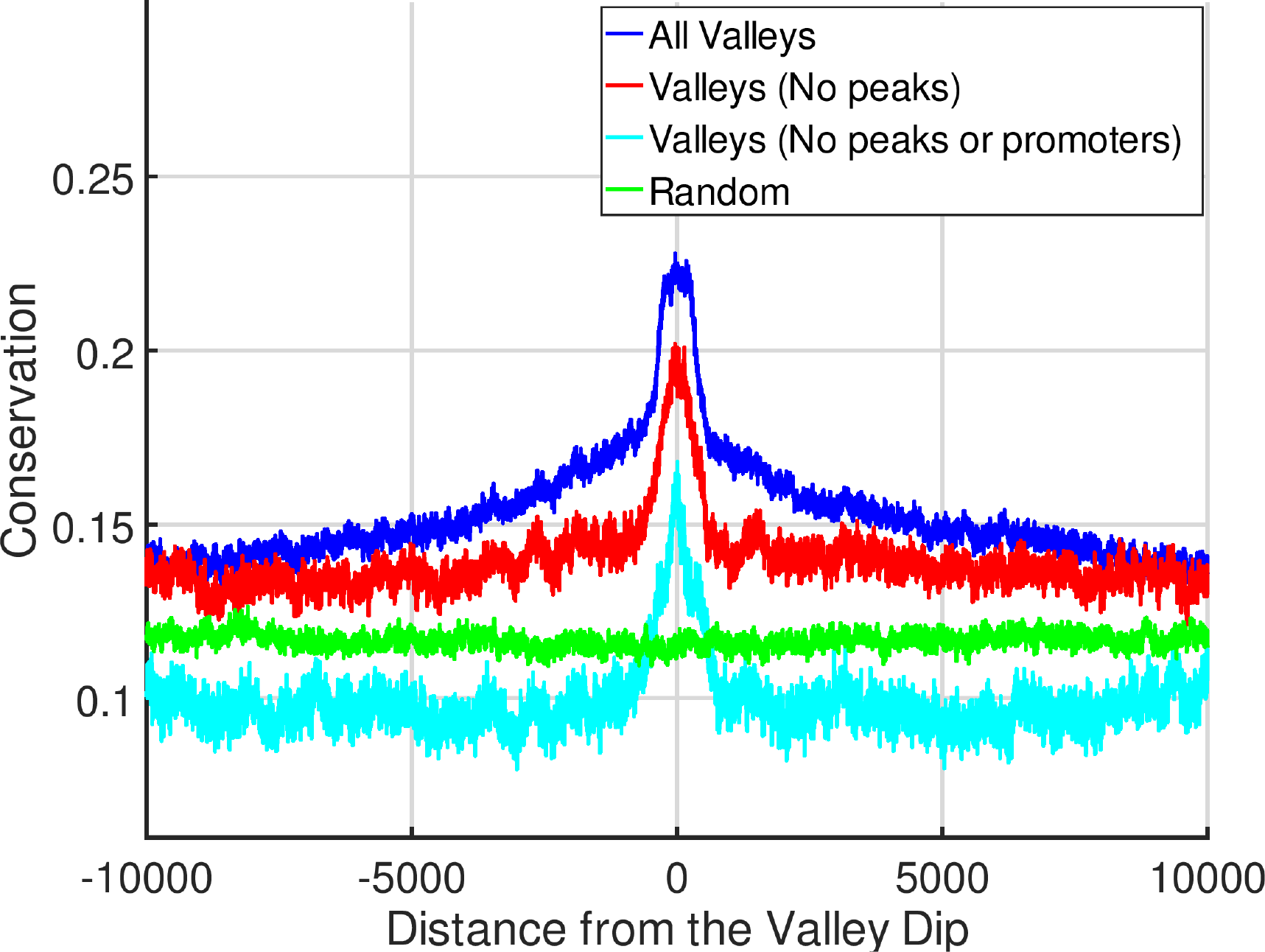
Transcription factor activity and conservation of H3K4me3 valleys. a) The number of overlaps between the top 1000 valleys and different transcription factors. X-axis shows the transcription factor and Y-axis shows the number of valleys that overlap with a peak of the corresponding transcription factor. The transcription factors are sorted with respect to decreasing number of valleys overlapping with them. b) Average conservation within 20,000 base pairs of the valley dips. X-axis shows the distance from the dip and y-axis shows the average PhyloP conservation score. The conservation around all valleys (blue), valleys that do not overlap with any H3K4me3 peaks (red), valleys that do not overlap with neither promoters and valleys (cyan), and randomized regions (green) are shown.

### Histone Valleys show High Conservation

We next computed the average conservation on the valleys identified by EpiSAFARI. To measure the conservation around the valleys, we aggregated the PhyloP conservation score (100 species genome-wide conservation obtained from The UCSC Genome Browser) around the 20,000 base pair vicinity of the reported dips of the valleys. Figure 3b shows the average signal around the dip. It is seen that there is a substantial increase in the average conservation signal around the valley dip and conservation decreases with increasing distance to the dip. It is expected that the H3K4me3 valleys will be enriched at the promoters, which already exhibit high conservation. To study whether the valleys that are outside any H3K4me3 signal enrichment show conservation, we computed the average conservation at the valleys that are do not overlap with H3K4me3 peaks (Fig. 3b). For this, we first called the peaks using MUSIC(14) and then excluded the valleys that overlap with any H3K4me3 peak. While the non-peak-overlapping valleys show slightly lower conservation, they still have higher conservation signal than random regions, where the random regions are generated by randomly shifting each valley within 1 megabase vicinity of itself. This result suggests that the valleys that are detected by EpiSAFARI that are outside the peak calls can reveal important functional information and bring new biological insight that may be missed by peak calls. We also computed the average conservation on the valleys that do not overlap with neither the H3K4me3 peaks nor with the promoters. These valleys represent the novel elements that EpiSAFARI identified. Figure 3b shows that the average conservation is high around the dips of the non-peak-non-promoter valleys. Overall, these results indicate that the valleys detected by EpiSAFARI may potentially contain genomic elements that are conserved among species.

### Nascent Transcription around the Valleys Correlates with Valleys’ Directionality

We also hypothesized that the valleys as detected by EpiSAFARI may contain cis-regulatory elements such as promoters and enhancers. One line of evidence for existence of these elements is the nascent transcription at the valleys. To study this, we used the global run-on sequencing (GRO-Seq) data for NA12878 sample (See Datasets). GRO-Seq data represent the genome-wide measurement of nascent transcription, i.e. RNA that has just been transcribed (or being transcribed) at each location in the genome(33). We aggregated the GRO-Seq signal from positive and negative strands around the 20,000 base pair vicinity of the reported dips of the valleys. Figure 4a shows the aggregation on different selections of valleys. When we look at all valleys, there is strong GRO-Seq signal pattern around the valleys compared to random regions in the genome. The positive and negative signals exhibit the general asymmetry observed in GRO-Seq signal profiles. We observed that the valleys that do not overlap with peaks show lower GRO-Seq signal but they contain higher GRO-Seq signal compared to the random regions (Fig. 4b). We also observed the double GRO-Seq peak pattern around the dips of the non-peak-overlapping valleys (Fig. 4b).

**Figure 4:**
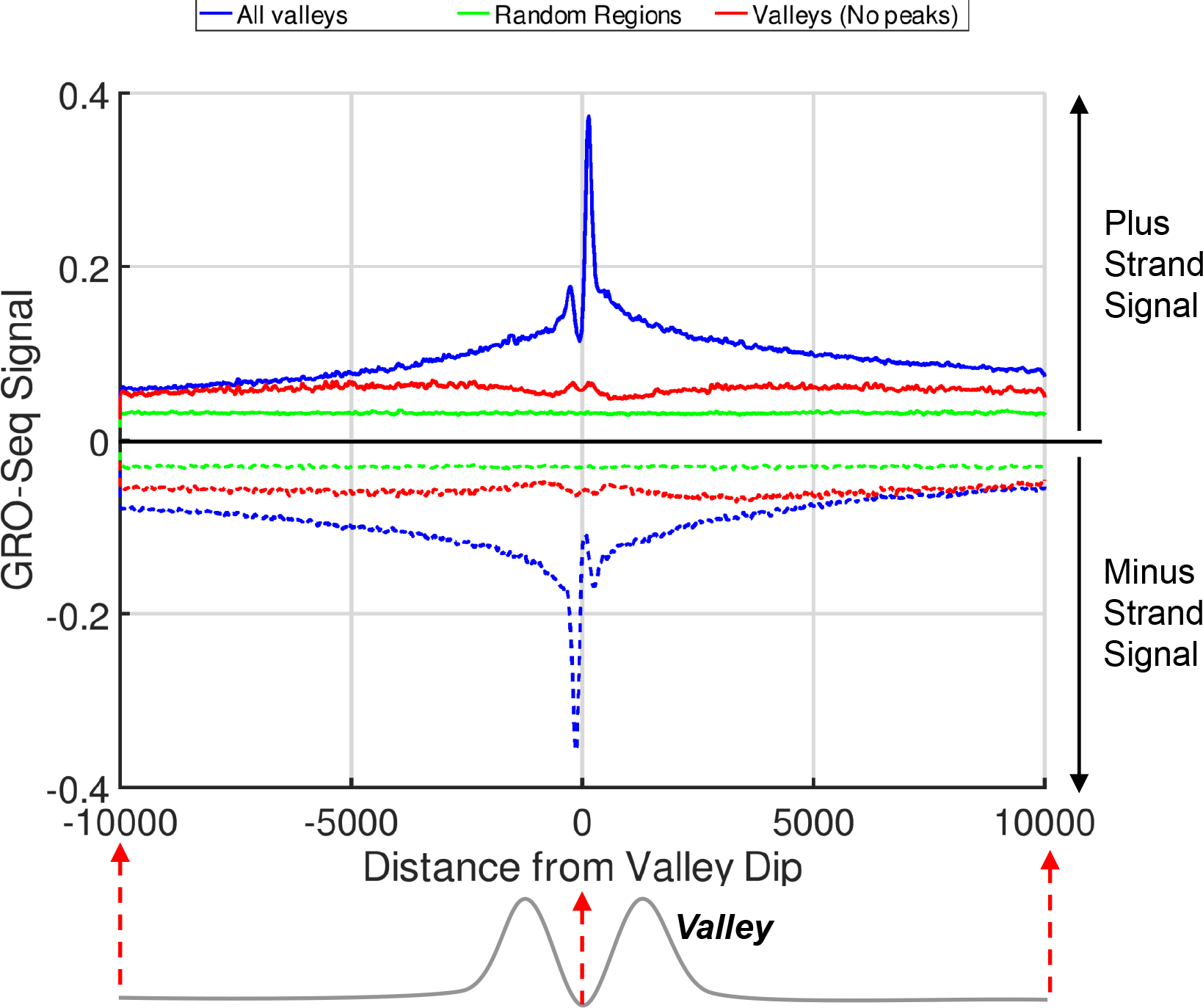
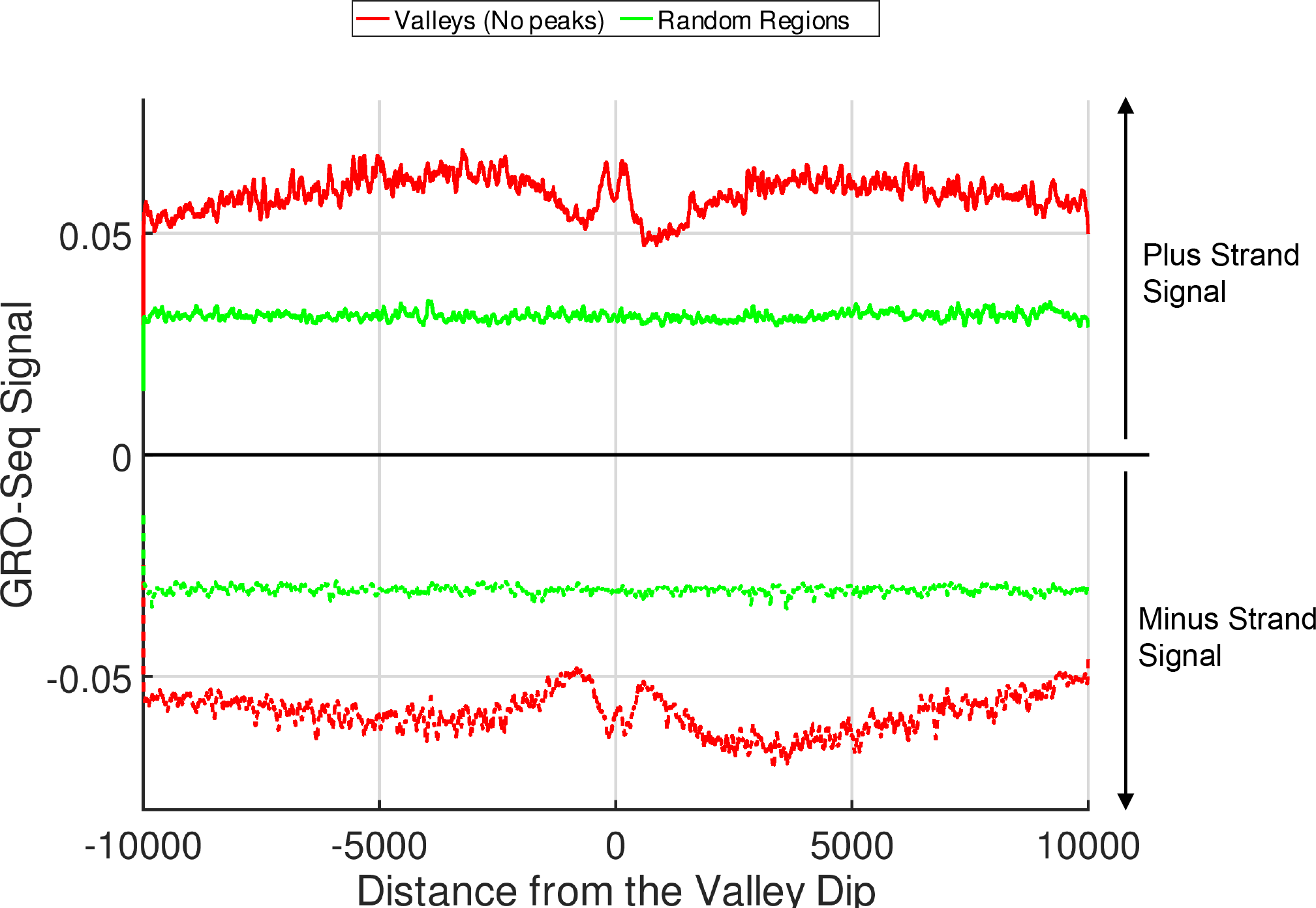
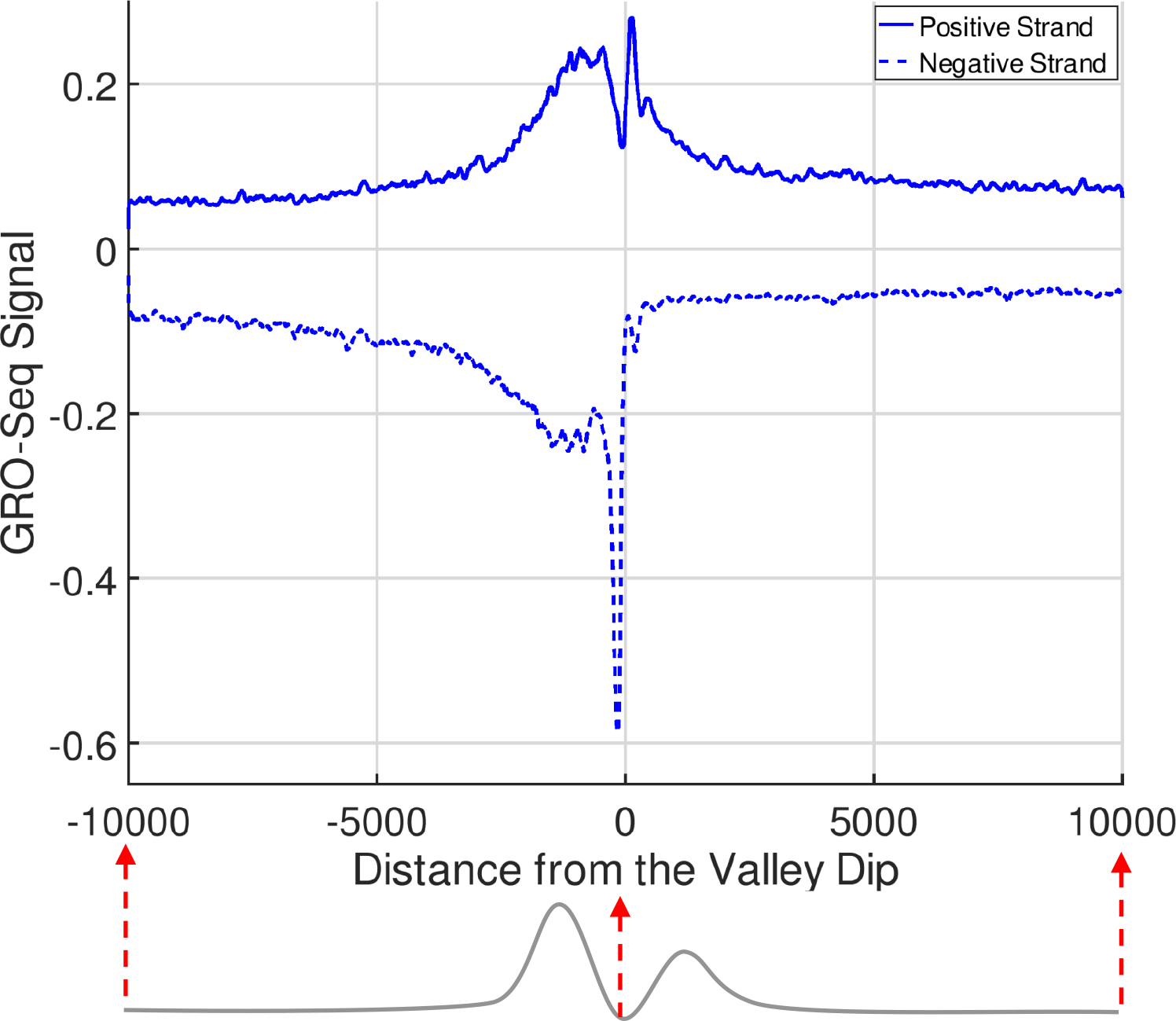
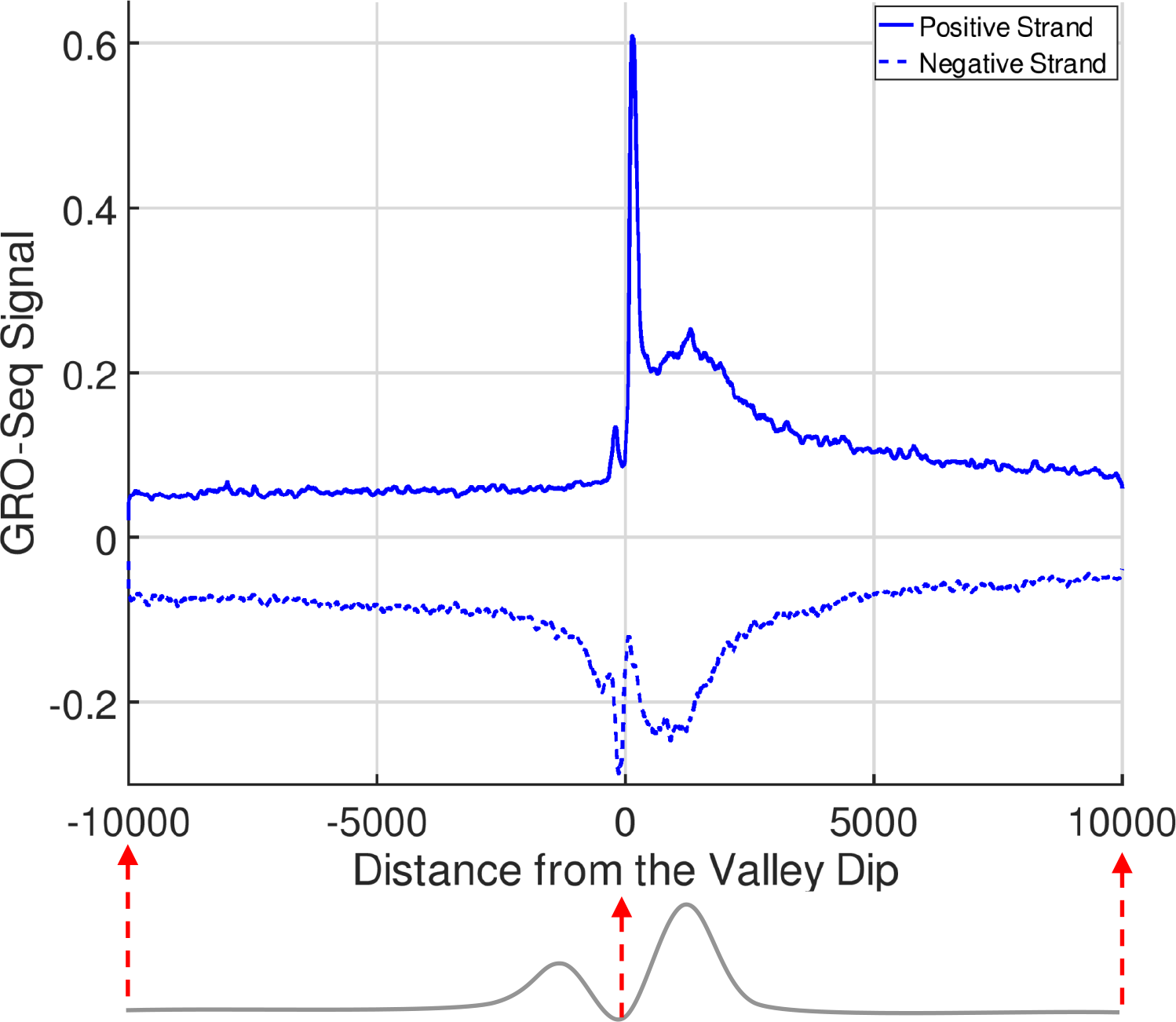
Analysis of nascent transcription around the H3K4me3 valleys. a) The aggregation of GRO-Seq signal within 20,000 base pairs of the dips reported by EpiSAFARI. All valleys (Blue), Randomized regions (Green), and valleys that do not overlap with any H3K4me3 peaks (Red) are plotted. Random regions are generated by randomly shifting the valleys within 1 megabase vicinity of the valley’s starting position. Plus and minus strand signals are plotted with straight and dashed lines, respectively. The valley below the figure aims to illustrate the valley’s positioning within 20 kilobase region. Two humps represent the two summits of the valley. Note that the summit locations in this illustration are not drawn to scale. The red dashed arrows indicate how the dip coordinates align with the x-axis of the aggregation plot. b) The aggregation of GRO-Seq signal for valleys that do not overlap with any peaks (Non-peak Valleys) and random regions are plotted to highlight the GRO-Seq signal on non-peak overlapping valleys. c) The aggregation of plus (straight line) and minus (dashed line) strand GRO-Seq signal within 20,000 base pairs of left-to-right valleys. The bottom illustration points out the fact that left summit is taller than right summit. d) The aggregation of GRO-Seq signal within 20,000 base pairs of right-to-left valleys where right summits are taller than left summits.

An important observation about the valleys is that valleys may show asymmetry with respect to the signal levels at the left and right summit positions. Several previous studies have highlighted the asymmetry of epigenetic signals and how this relates to transcriptional activity(34). We evaluated whether the asymmetry of the valley shapes detected by EpiSAFARI gives useful information about the nascent transcriptional activity. In order to study the relation between valley shape asymmetry and transcriptional activity around the valleys, we divided the valleys into two groups based on the signal levels at the summits of each valley. First group of valleys, we call left-to-right valleys (Bottom illustration in Fig. 4c), have higher signal on the left summit compared to right summit. Second group, we named right-to-left valleys, (Bottom illustration in Fig. 4d) contains higher signal on the right summit compared to the left summit. We then computed the average GRO-Seq signal profile around the dips of the valleys. Figure 4c and 4d show the average GRO-Seq signal around the 20,000 base pairs vicinity of the valley dips for left-to-right and right-to-left valleys, respectively. For left-to-right valleys, there is a sharp peak on the negative strand signal to the left of the dip position. The positive signal, while still high, does not show a corresponding sharp peak. In other words, left-to-right valleys are enriched in terms of negatively oriented nascent transcription. Similar pattern is seen for the right-to-left valleys (Fig. 4d) albeit on the positive strand. The positive nascent transcription is highly enriched with a very sharp peak on the positive strand GRO-Seq signal. While there is some transcription on negative strand, the sharp peak does not exist. Overall, these results provide supporting evidence that the valleys detected by EpiSAFARI contain genomic elements of potential functional role and the directionality of transcriptional activity around them.

### Methyl-Valleys are Conserved and are Enriched in Chromatin Structure Associated Transcription Factor Binding

As another application of EpiSAFARI, we next focused on analysis of the valleys in the DNA methylation signal of H1 embryonic stem cell line (H1hESC). The valleys in DNA methylation signals have been shown to contribute to important biological phenomena(15, 35). The current standard method for measuring genome-wide DNA methylation levels is whole genome bisulfite sequencing (WGBS). One important aspect to consider while analyzing this data is that the WGBS signal is sparse because it is measured only at the CpG nucleotides. This makes valley detection from WGBS data challenging. To get around this issue, EpiSAFARI utilizes a sparse mode to smooth the sparse signal profiles in the spline fitting step. In the sparse mode, EpiSAFARI does not use genomic locations for which signal is exactly 0. This way, EpiSAFARI interpolates the sparse signal using only at the locations that it is measured. We first downloaded the processed the WGBS signal profile from Roadmap Epigenome Project(36). This signal profile measures the fraction of methylated versus non-methylated cytosine residues at the CpG di-nucleotides. We identified valleys in DNA methylation signal, i.e. methyl-valleys, using sparse mode of EpiSAFARI. In valley detection, we set *l_min_* = 250, *l_max_* = 2,000 and excluded the valleys that contain less than 20 CpG di-nucleotides as these may correspond to valleys with very sparse signals. While smoothing DNA methylation signals, *l_w_* = 5,000 is used.

To evaluate the interspecies conservation on methyl-valleys, we computed the average PhyloP conservation score(10) within the methyl-valleys. We aggregated the conservation signal around 20,000 base pairs vicinity of the methyl-valley dip as reported by EpiSAFARI. Figure 5a shows the average signal for all valleys and the valleys that do not overlap with promoters. Compared to the random regions, the valleys show a higher conservation signal. We next analyzed whether the methyl-valleys are enriched in transcription factor binding. We overlapped the valleys with the transcription factor ChIP-Seq peaks for H1hESC from the ENCODE Project. Figure 5b shows the fraction of top methyl-valleys that overlap with a transcription factor binding peak. The top methyl-valleys are highly enriched (higher than 90%) in terms of transcription factor binding. We then excluded the methyl-valleys that overlap with promoters and computed the fraction of non-promoter-overlapping methyl-valleys that overlap with transcription factor binding peaks (Fig. 5b). Top valleys in this category are also highly enriched in transcription factor binding. We next computed the fraction of top 1000 valleys that overlap with each transcription factor. Specifically, we overlapped the peaks for each of the transcription factor with the top 1000 methyl-valleys. Figure 5c shows the fraction of top methyl-valleys that overlap with the binding peaks of each transcription factor in decreasing order. Large fraction of methyl-valleys overlaps with transcription factors such as CTCF (60%), Rad21 (50%), and Znf143 (50%), which are known regulators of three-dimensional chromatin structure. To study the co-binding preference of the transcription factors, we computed which transcription factors are bound to each of the top methyl-valleys. We computed a binary matrix whose rows are methyl-valleys and columns are transcription factors. Each entry in the matrix represents whether there is binding of the transcription factor in the column to the corresponding methyl-valley. We next clustered the matrix for visualization (Fig. 5d). The chromatin structure associated factors predominantly co-associate with each other around the methyl-valleys. Moreover, the methyl-valleys that do not overlap with promoters are also enriched in binding of Rad21 and CTCF transcription factors (Fig. 5e). We also saw that a similar result has been also found in another study(37) where the authors show that chromatin structure associated transcription factors are enriched in undermethylated regions.

**Figure 5:**
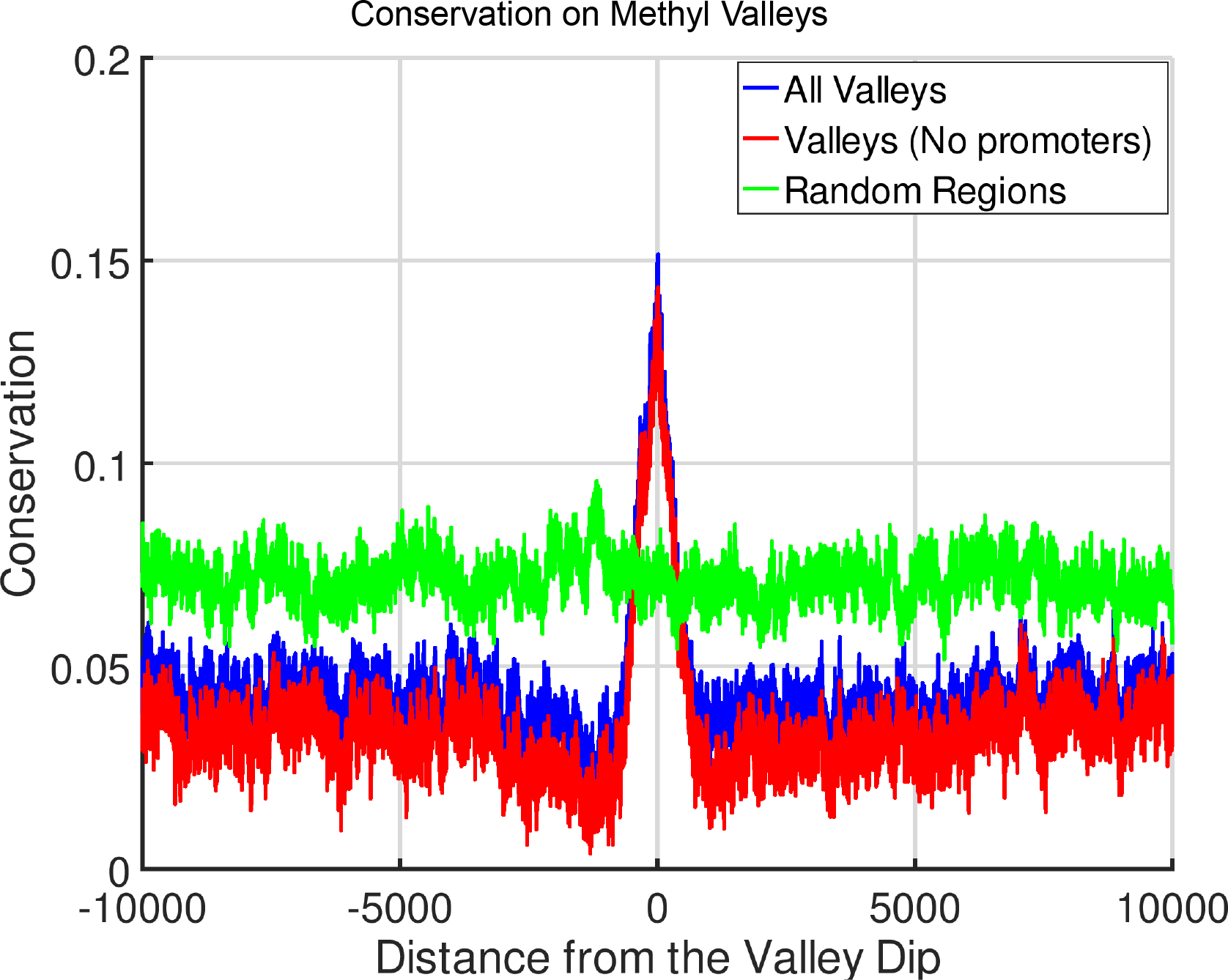
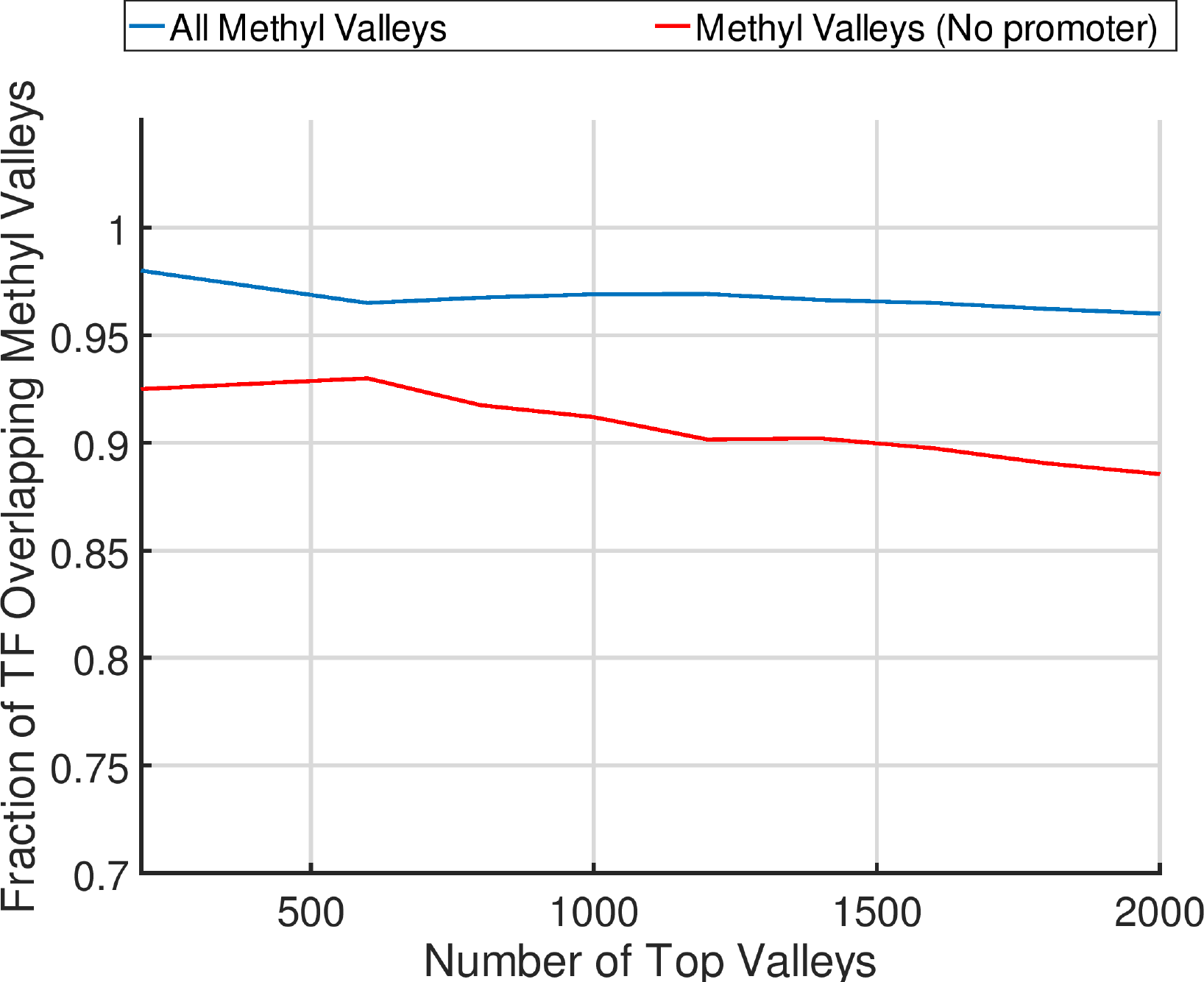
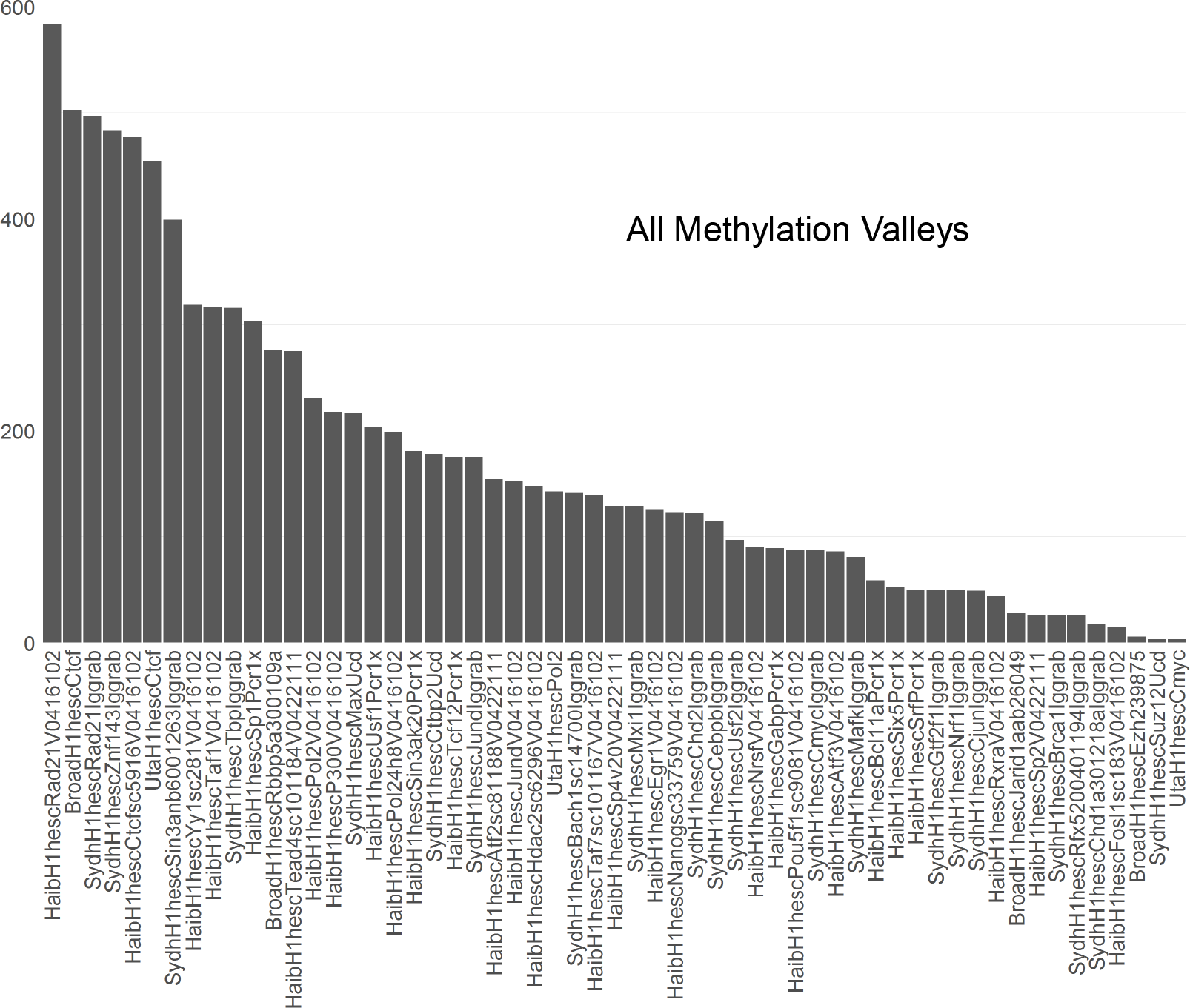
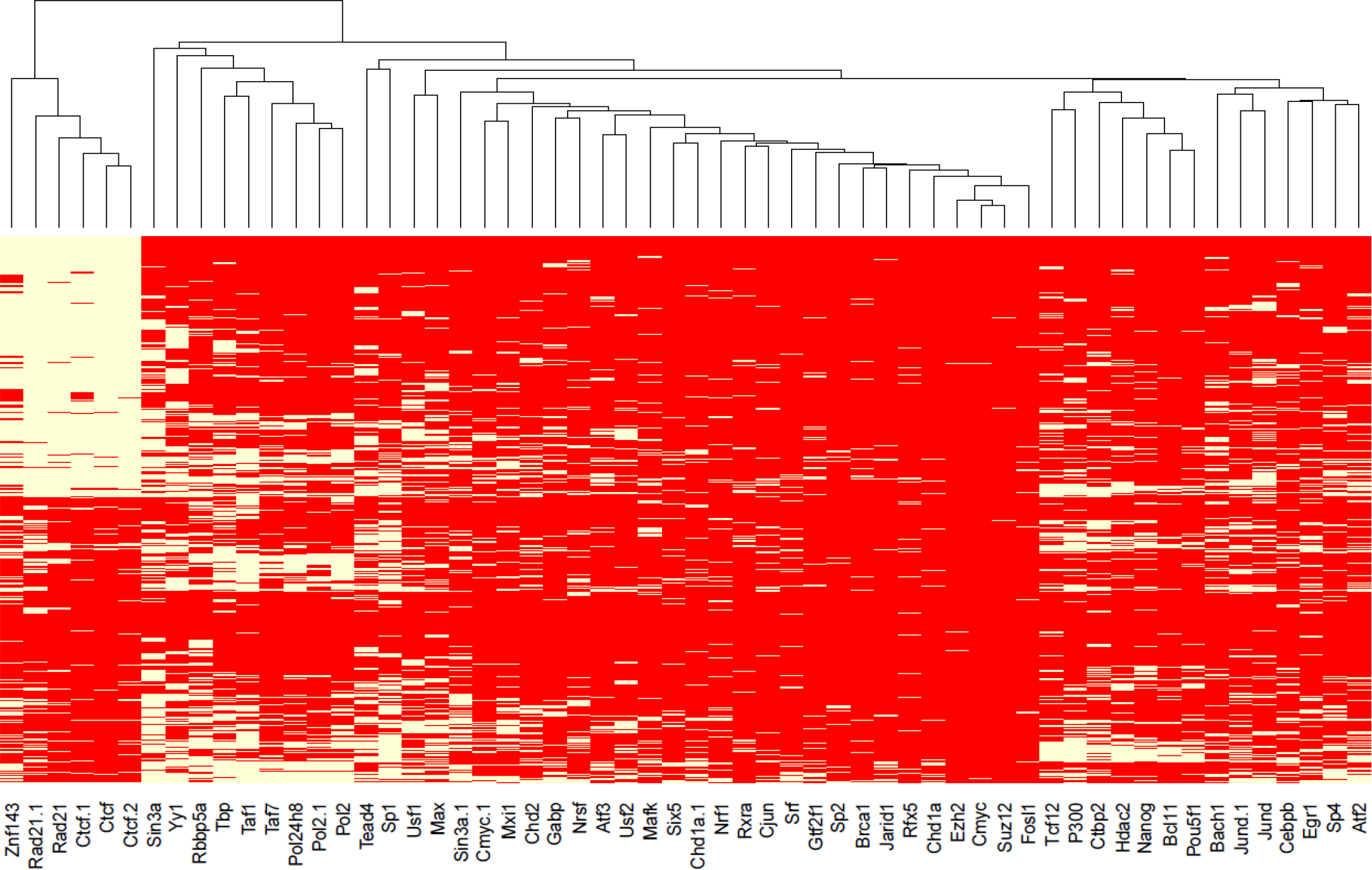
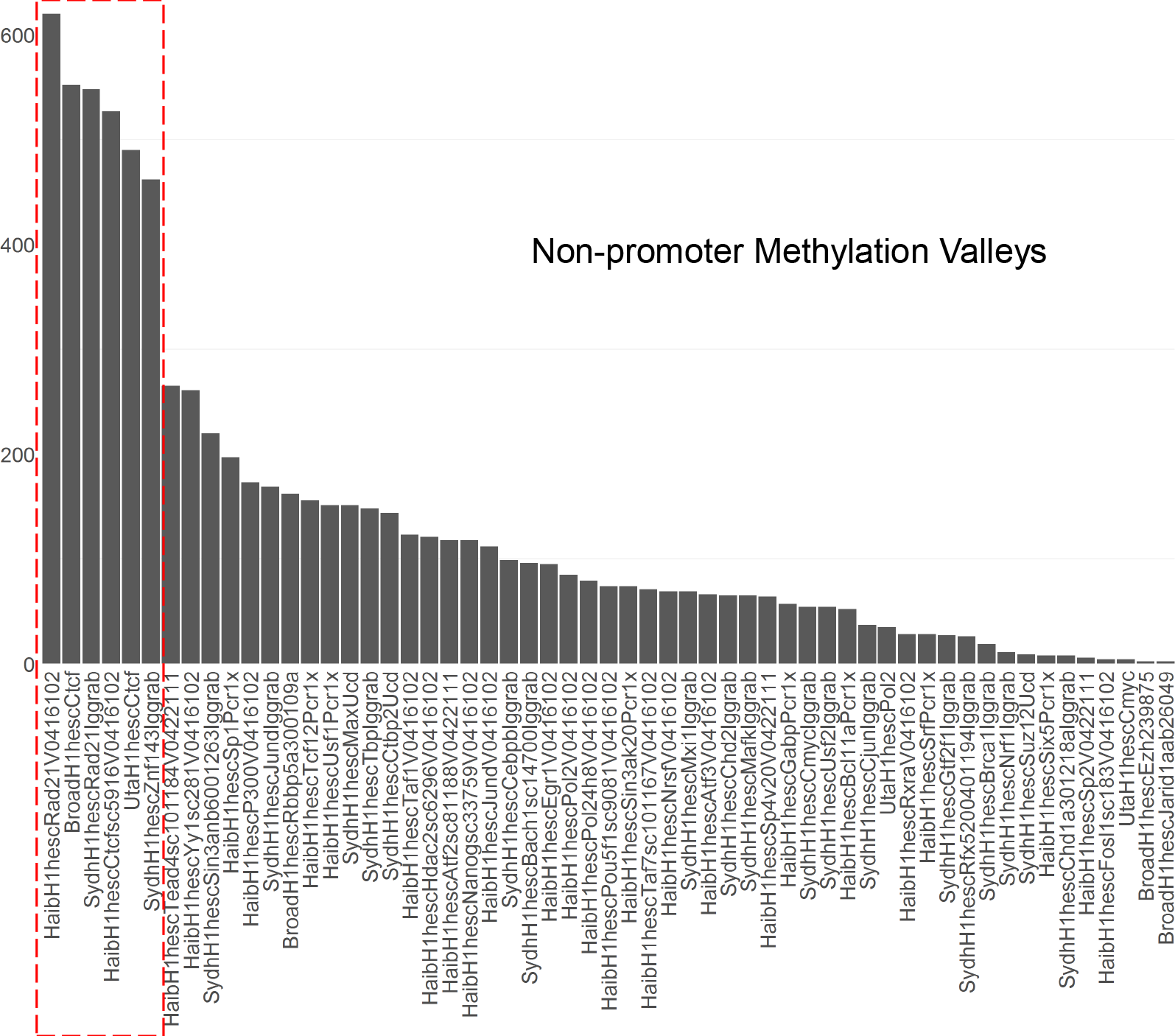
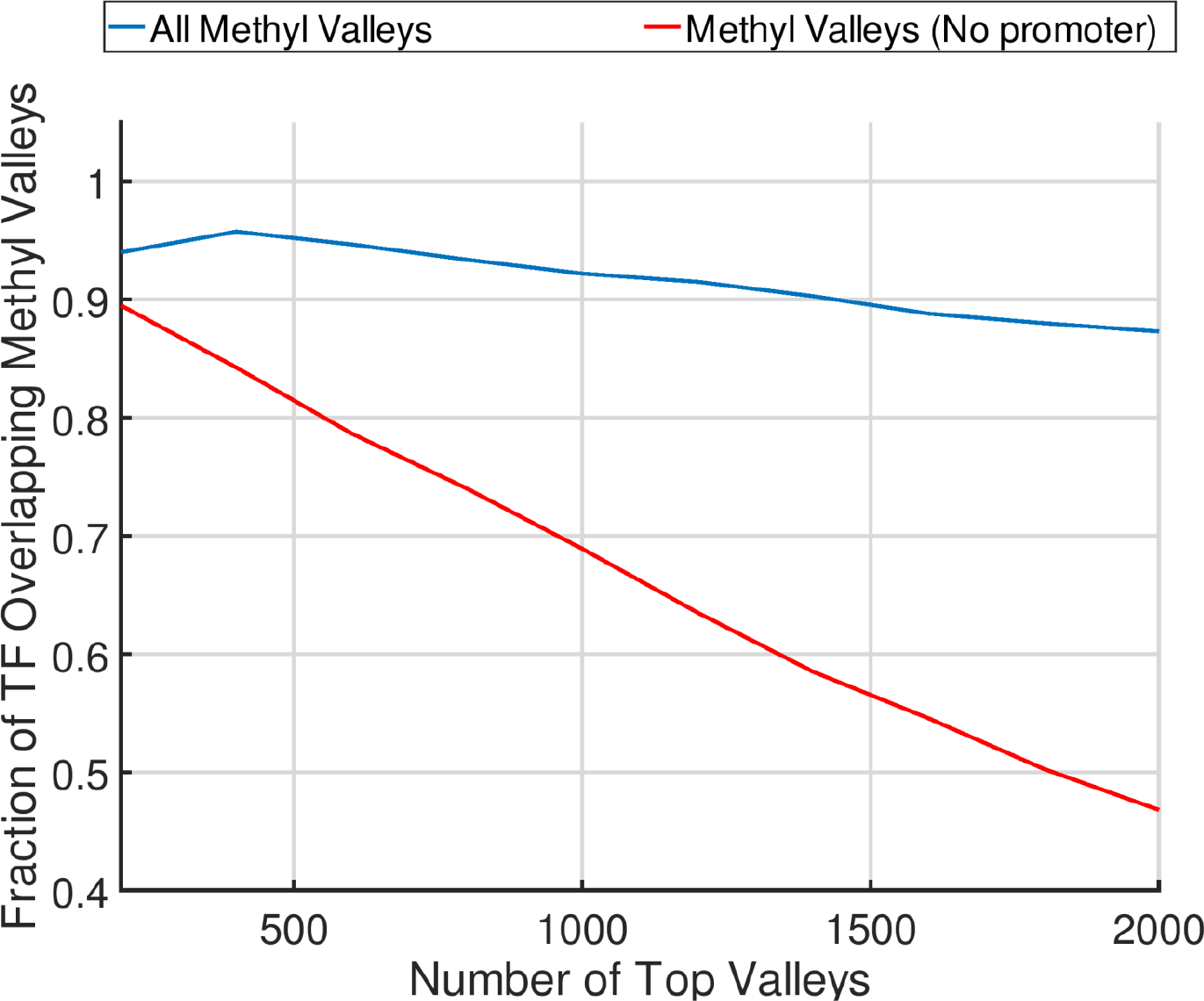
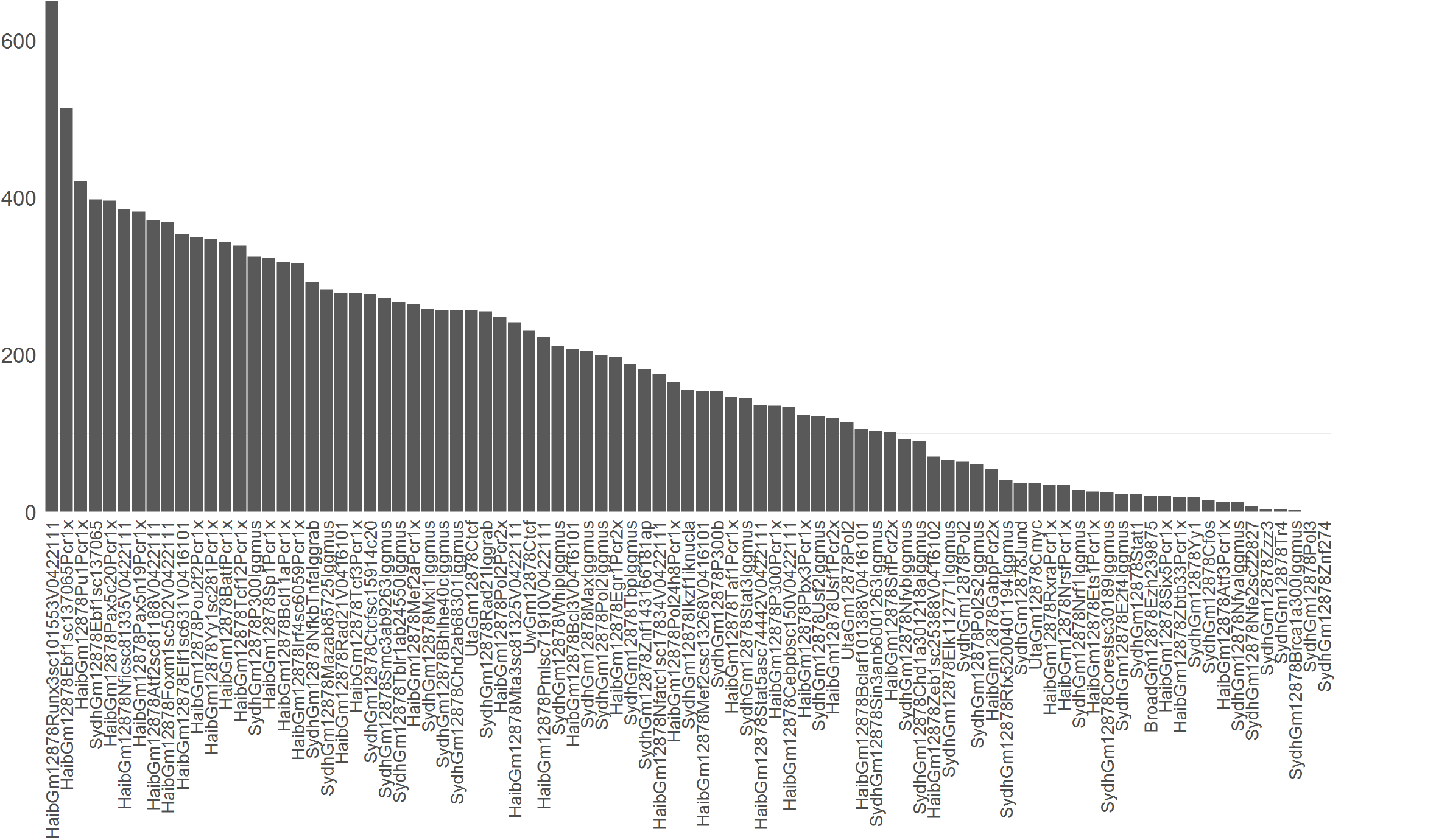
Characterization of the methyl-valleys. a) Conservation around the 20,000 base pair vicinity of the dips of methyl-valleys as reported by EpiSAFARI. All valleys (blue), valleys that do not overlap with promoters (red), and random regions (green) are shown. b) The fraction of the top 2000 methyl-valleys that overlap with a H1hESC transcription factor binding peaks. X-axis shows the top number of valleys and y-axis shows the fraction of valleys that overlap with at least one transcription factor peak. Fractions for All methyl valleys (blue) and methl-valleys that do not overlap with any promoters (red) are illustrated. c) Overlap of top 1000 methyl-valleys with transcription factors. The bars are sorted with respect to decreasing number of overlaps. The transcription factors are shown on x-axis and the number of overlapping methyl-valleys are shown on y-axis. d) Heatmap of the transcription factor binding on the top 1000 methyl-valleys. Each row is a methyl-valley and each column is a transcription factor. Within a row, a white color indicates an overlap with the valley corresponding to the row and the peaks of the transcription factor that is corresponding to the column. Red indicates no overlap between the transcription factor’s peaks and the valley. The rows and columns are sorted by hierarchical clustering. e) Overlap of top 1000 non-promoter associated methyl-valleys with transcription factors. The top transcription factors associated with chromatin structure are highlighted with a dashed red rectangle. f) The fraction of the top 2000 methyl-valleys in GM12878 that overlap with a transcription factor peak. g) The frequency of transcription factors that overlap within top GM12878 methyl-valleys.

We also used EpiSAFARI to detect the methyl-valleys for NA12878 cell line using a publicly available dataset. While we did observe that the top methyl-valleys are enriched in transcription factor binding (Fig. 5f), we did not observe the enrichment of the chromatin structure associated transcription factors (Fig 5g).

### Supervalleys are Enriched at Promoters of Developmental Genes in Embryonic Stem Cell Line

An interesting question about the valleys is related to whether the clusters of H3K4me3 valleys, i.e., valleys that are close to each other reveal important biological insight. The motivation behind this idea is that valleys that are clustered together in genomic coordinates may represent super-enhancer regions(38), which have been shown to associated with important biological phenomena such as cancer initiation. In addition, several studies have shown that the broad H3K4me3 peaks are enriched around genes with certain biological roles(39, 40). We use the number of valleys that make up the signal at the promoters to measure the signal breadth. We overlapped the H3K4me3 valleys for H1HESC cell line with the promoters and identified the promoters with large number of valleys around them (Fig. 6a). We refer to these large clusters of valleys as supervalleys. We sorted the promoters with respect to decreasing number of overlapping valleys and selected the top 200 genes with the largest number of valleys around their promoters. We next performed GO enrichment analysis on these genes. Interestingly, these genes have significant enrichment of biological processes related to development, differentiation, and DNA binding (Fig 6b). Consequently, the clustering of the valleys with respect to proximity in genomic coordinates may be an indicator of elements with important biological functions. When we performed the clustered valley analysis using H3K4me3 valleys of NA12878 cell line, we did not find any significant functional category.

**Figure 6:**
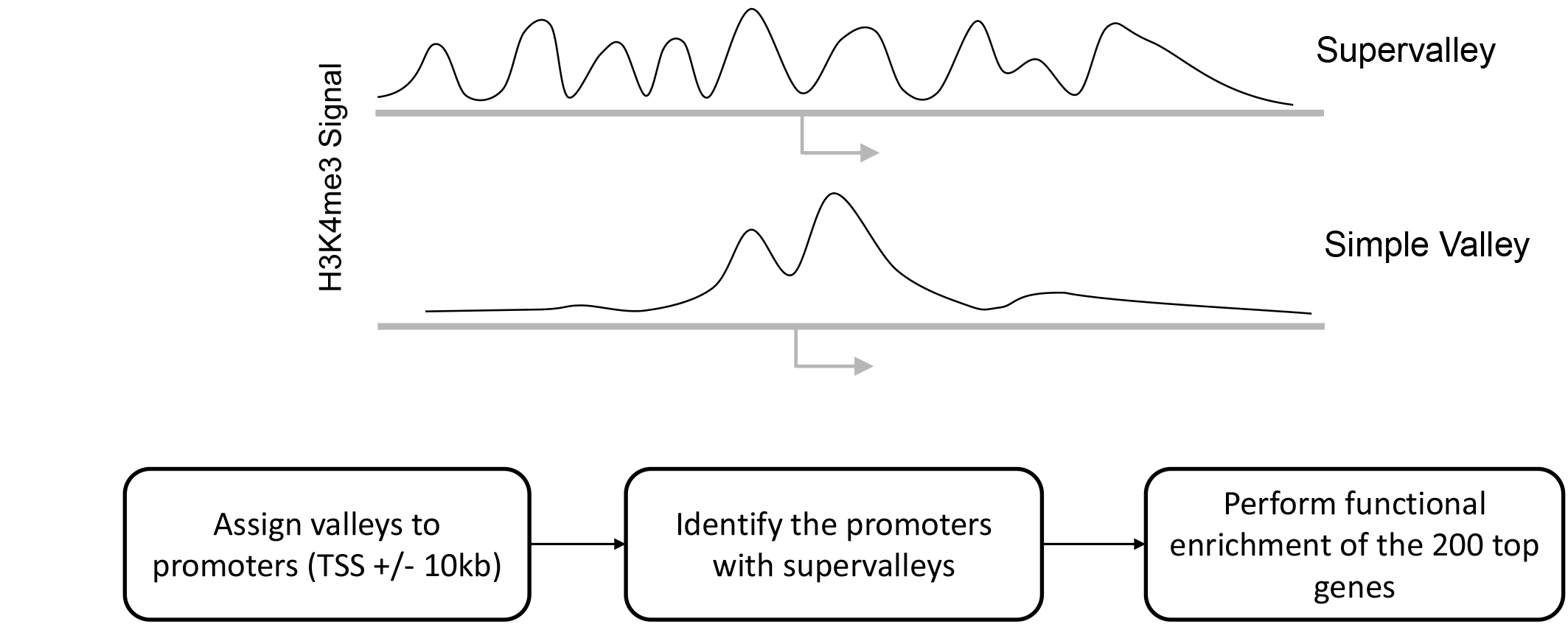
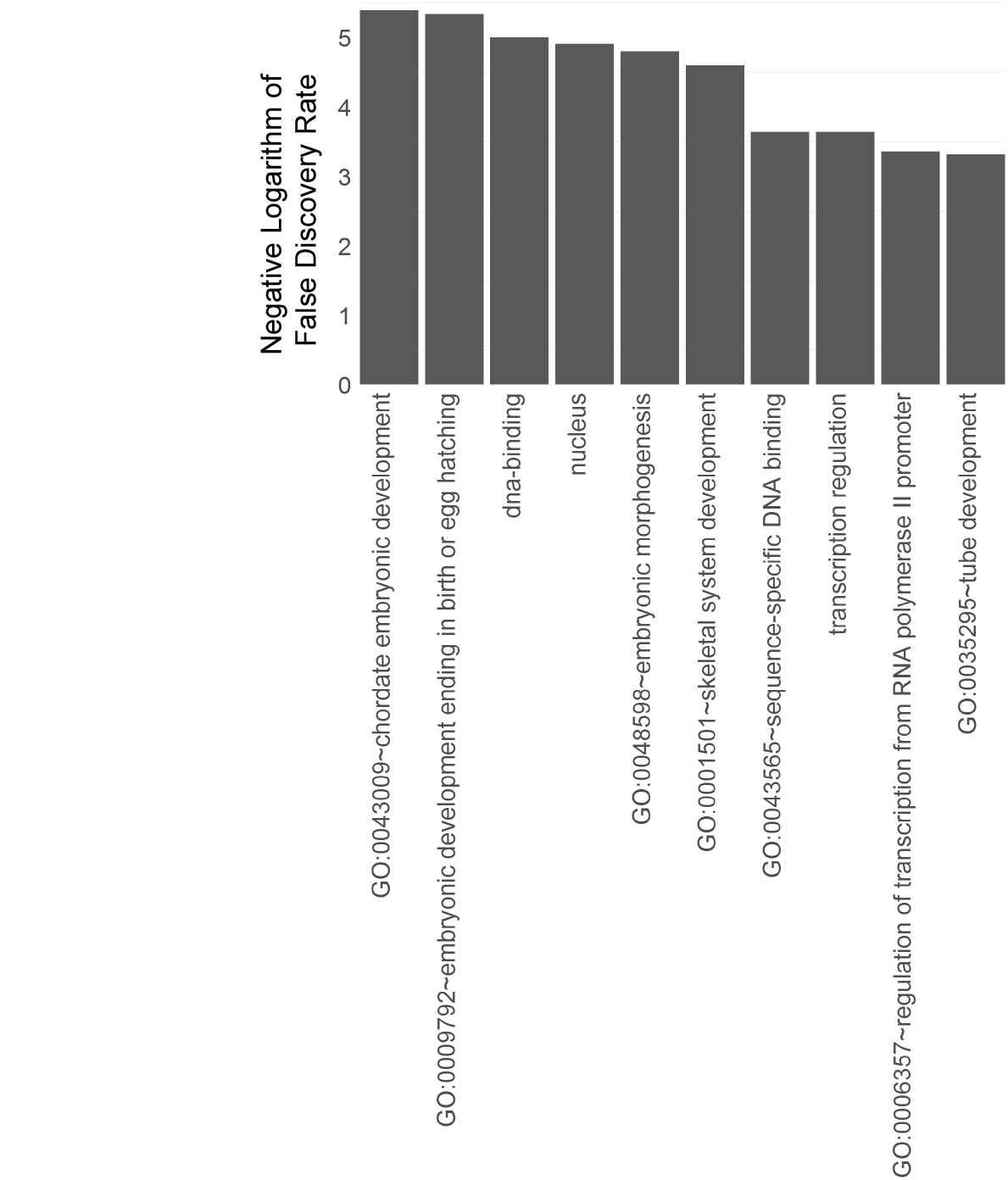
GO enrichment of the genes whose promoters overlap with multiple valleys, i.e. supervalleys. a) Illustration of a simple valley and a H3K4me3 supervalley. The simple valley consists of a double-peak pattern with a valley in it. The supervalleys contain many consecutive valleys that are clustered within a small genomie distance. Block diagram below illustrates the detection of supervalleys on promoters. The valleys are assigned to gene promoters such that a promoter is defined as 10kb vicinity of the transcription start site (TSS). The genes whose promoters overlap with the largest number of valleys are identified. The top 200 genes are used in gene ontology enrichment. b) The most significant 10 GO terms (x-axis) that are detected from ontology enrichment as sorted with respect to enrichment false discovery rate (y-axis)

## Discussion

EpiSAFARI is a novel statistical method for detecting the valleys in the fluctuations of epigenetic signal profiles. The valleys represent important biological events such as transcription factor binding sites, open chromatin, and cis-regulatory elements such as enhancers. EpiSAFARI summarizes the signal profile data and generates several useful outputs. First is the set of detected valleys and metrics (such as signal levels, mappability, and nucleotide composition) that can be used to filter the valleys. Second, the smoothed signal profiles can be useful for visualizing and other downstream analysis. EpiSAFARI uses a smoothing approach which does not make assumptions on the underlying experimental assay. This is advantageous because it enables analysis of any type of genome-wide signal profile such as the sparse profiles (such as DNA methylation). EpiSAFARI represents a parallel approach to complement the current epigenetic data analysis methods that predominantly rely on detection of broad regions, e.g. peaks and differentially methylated regions. While the peaks represent biologically important genomic events and elements, valleys help refine the peaks such that the detailed information in the signal fluctuations are utilized for delineation of the peak regions.

One of the main challenges related to the valley-centric analysis of epigenetic data is concretizing the definition “valley calling” process. EpiSAFARI treats the valleys to be dips in the signal that are between two summits but this definition could potentially be revised to ensure that the valleys represent the functionally most meaningful regions. Another limitation that we have faced is defining quality metrics for the detected valleys. The hill score aims to measure the valley quality but we observed that it may be affected to a certain extent by the signal smoothing parameters. More robust measures of valley quality can elucidate the valley quality. Another challenge is defining statistical models for valley calling. Although we evaluated several statistical models that evaluate the significance of the valleys, the definition of statistical significance of valleys should be studied in more detail.

Several previous methods have utilized valleys in different contexts. These methods rely on smoothing of signal using kernel-based approaches (such as Gaussian(24) and wavelet filtering(19)) or modeling of the read clusters (such as PARE). In comparison, EpiSAFARI is advantageous to these methods for two reasons. Firstly, EpiSAFARI incorporates mappability and nucleotide content in filtering of the detected valleys. As we have demonstrated, these factors may create false positive valleys. Secondly, the kernel smoothing-based methods may fail to smooth sparse signals (such as DNA methylation) because smoothing of sparse signals will introduce many false positive valleys. On the other hand, EpiSAFARI computes an interpolation of the sparse signal to efficiently build a continuous smoothing of the sparse signals. Thus, the spline-based modeling of EpiSAFARI separates it from previous methods for modeling of both continuous and sparse signal profiles.

In the analysis of H3K4me3 valleys, the valleys outside the identified peaks show significant conservation. This highlights the potential novel utility that the valley-based analysis of the signal profiles can provide. As most of the ChIP-Seq analysis pipelines are focused on peak calls, the valleys can provide new insight into characterizing ChIP-Seq data more comprehensively than focusing just on the peak calls. In addition, the newly developed assays such as ATAC-Seq(41) can benefit greatly by the information provided by EpiSAFARI. In this respect, the features generated by EpiSAFARI can complement the existing peak calling methods for the new assays. In summary, EpiSAFARI represents an efficient tool that can be integrated directly into the functional genomics analysis pipelines.

### Data Availability and Accession Numbers

The H3K4me3 histone modification ChIP-Seq and DNase data, and transcript expression quantifications for K562, GM12878, and H1hESC cell lines are downloaded from ENCODE project website (http://hgdownload.soe.ucsc.edu/goldenPath/hg19/encodeDCC). The transcription factor binding peaks for K562, GM12878 and H1hESC cell lines are downloaded from the uniformly processed datasets of the ENCODE Project. The conservation scores are downloaded from the 100-way PhyloP track of UCSC Genome Browser. Whole genome bisulfite sequencing-based DNA methylation data for H1hESC data is downloaded from the Roadmap Epigenome Project Data Browser. GRO-Seq data is downloaded from GEO website with accession number GSM1480326. The GO enrichment analysis is performed using DAVID website(42). The random valleys in aggregation analyses and plots are generated by randomly shifting each valley within 1 megabase vicinity of itself. The whole genome bisulfite sequencing-based DNA methylation data for GM12878 cell line is downloaded from GEO web site with accession number GSM2772524.

The multi-mappability profile is obtained as described in a previous publication(14). In summary, the genomes are fragmented into fragments of the desired read length (denote by *l_read_*) and these are mapped back to the reference genome by allowing multimapping reads. After the reads are mapped, we count the number of reads that are mapping to every genomic position. For any genomic position that is uniquely mappable, this computation yields exactly 2 × *l_read_* at that position. For any genomic position that is multi-mapped, the number of overlapping reads at the position will be higher than 2 × *l_read_*, therefore we call this profile the multi-mappability profile. This profile quantifies the multi-mappability of each position in the genome for the given read length of *l_read_*.

## Supplementary Data

Supplementary Data are available upon request.

## Acknowledgements

None.

## Supplementary Figures

**Supplementary Figure 1.**
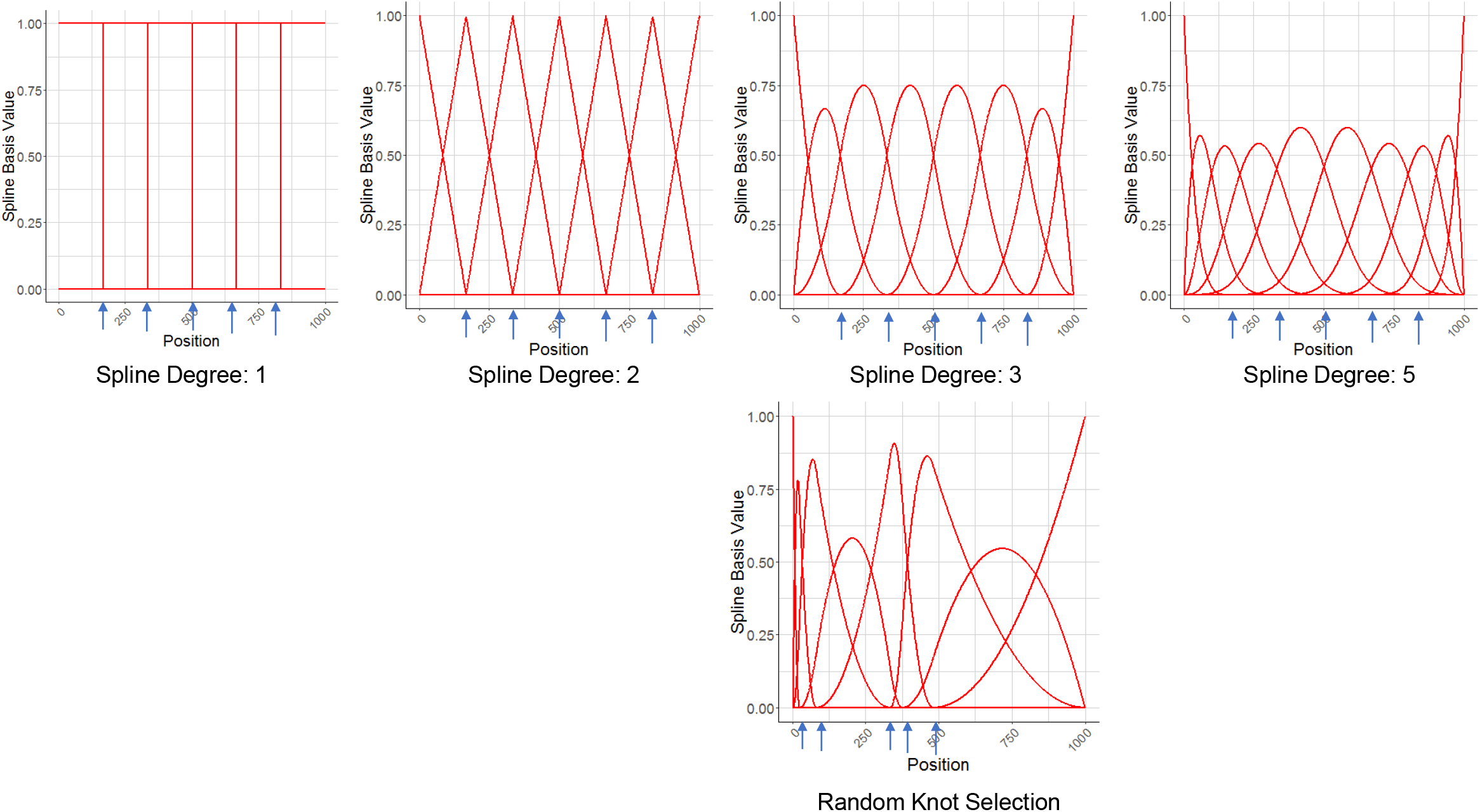

**Supplementary Figure 2.**
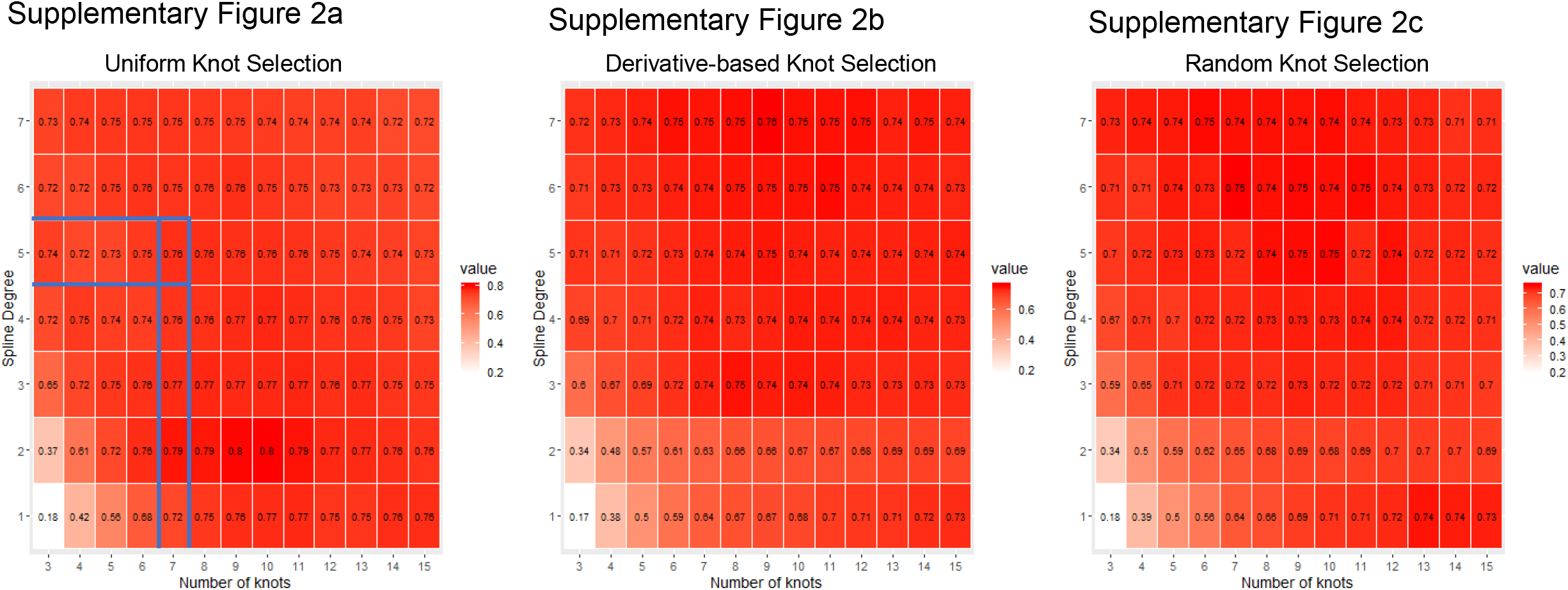
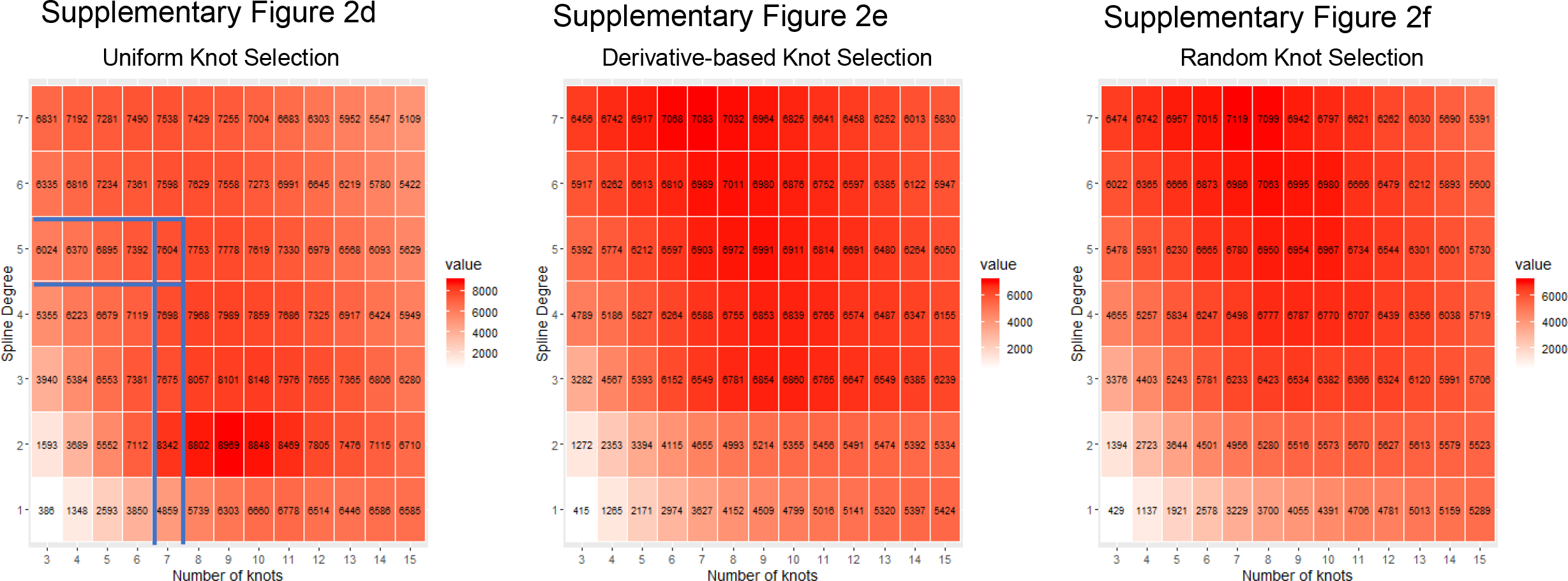
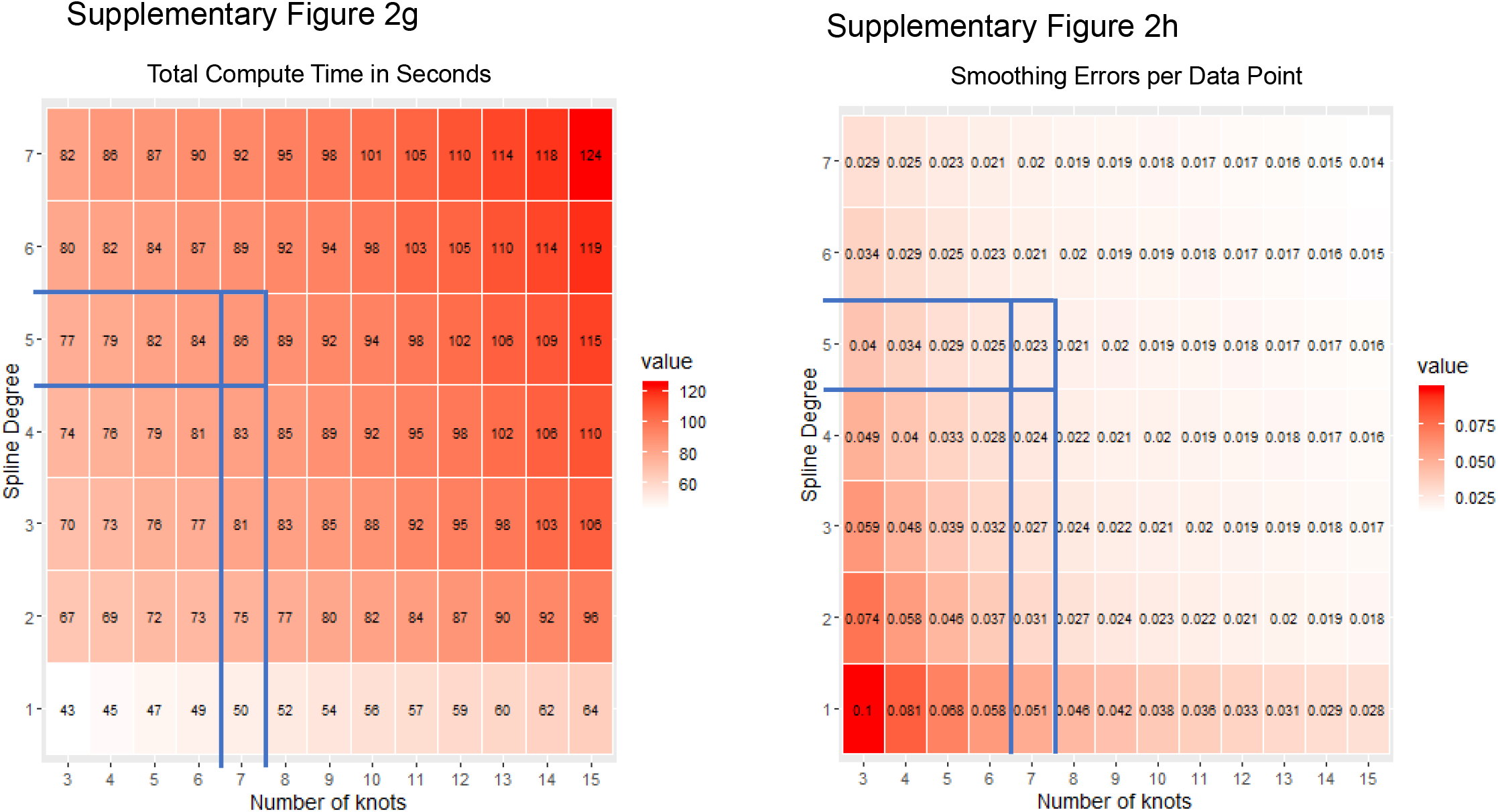

**Supplementary Figure 3.**
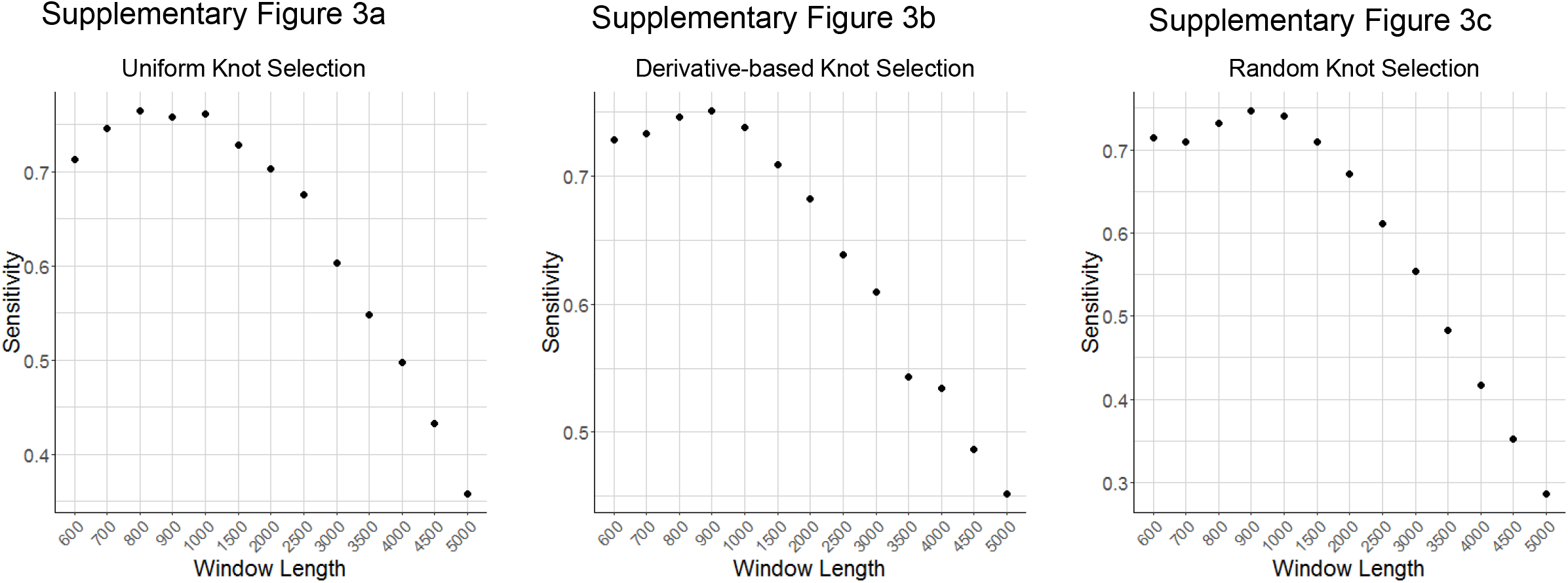

**Supplementary Figure 4.**
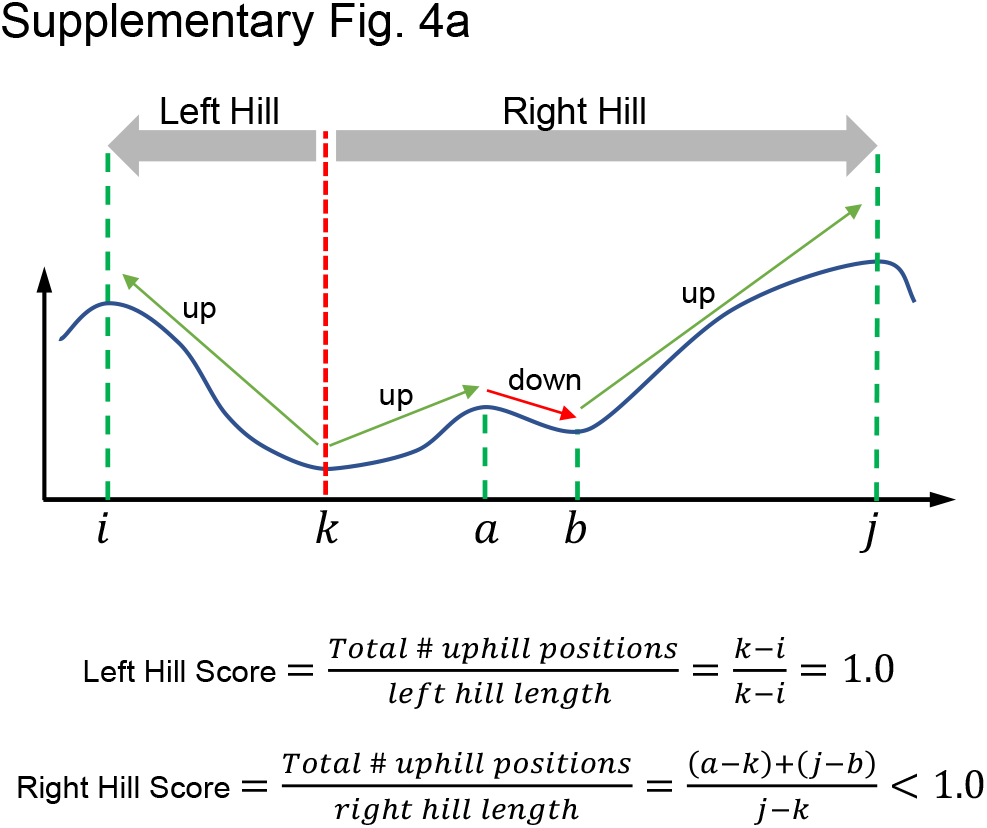
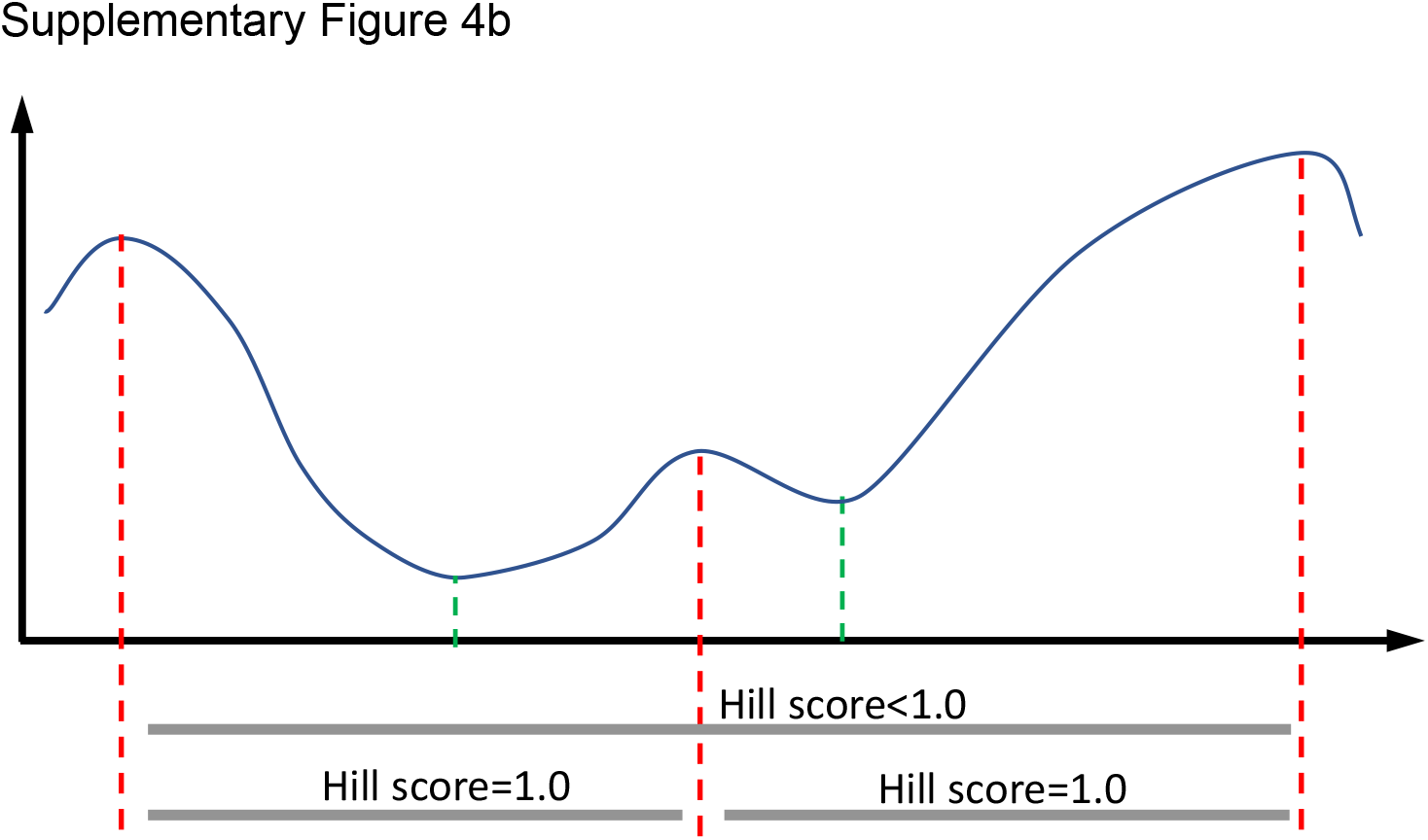
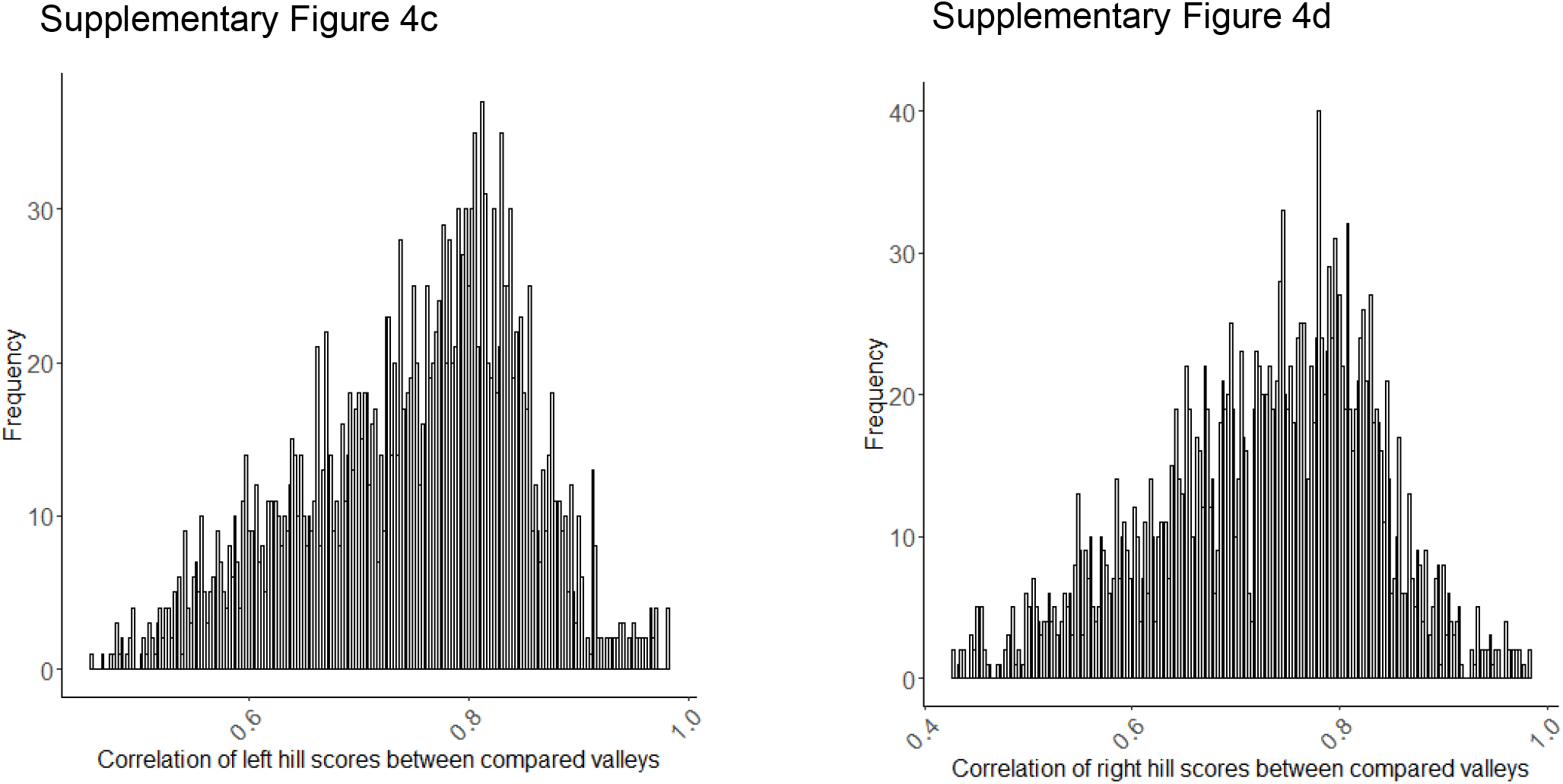
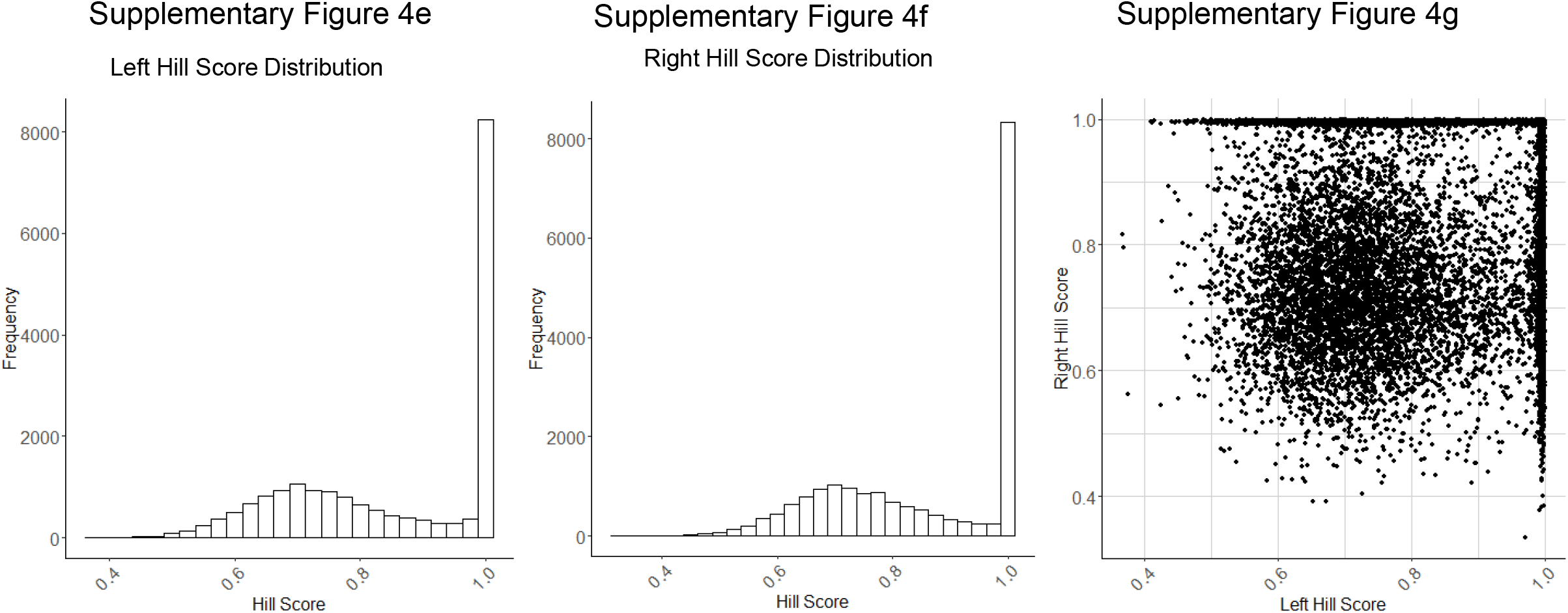
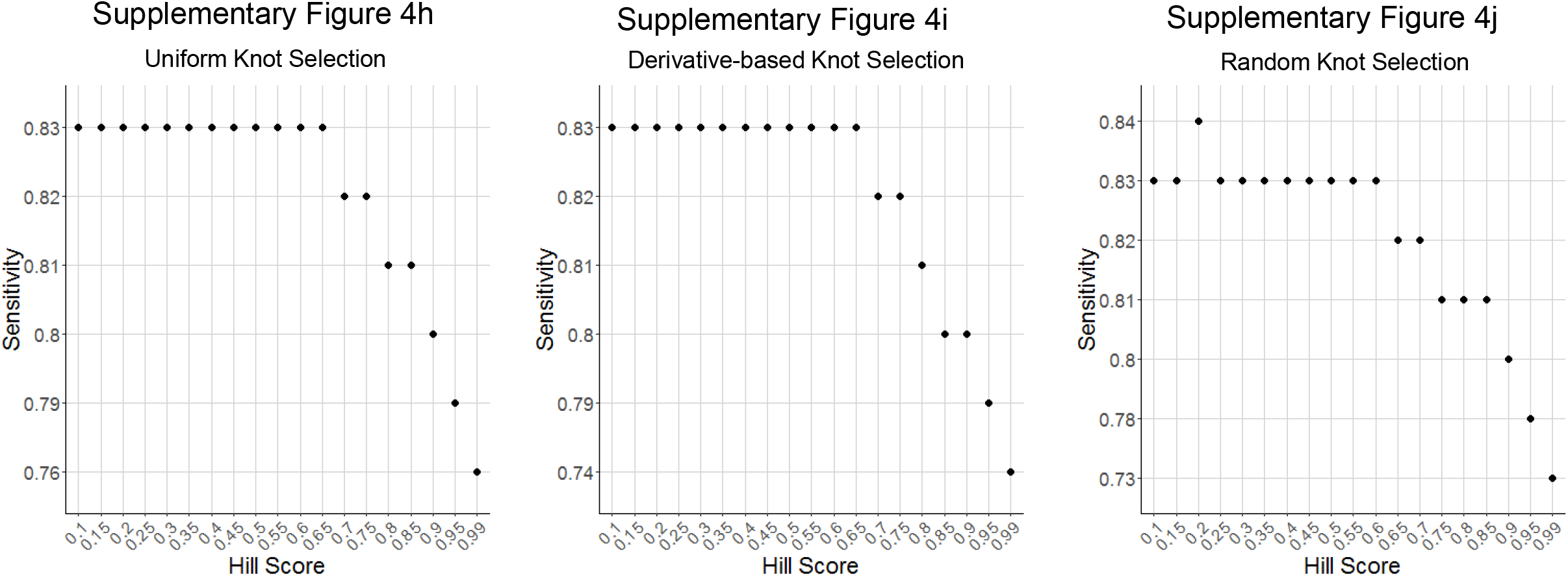
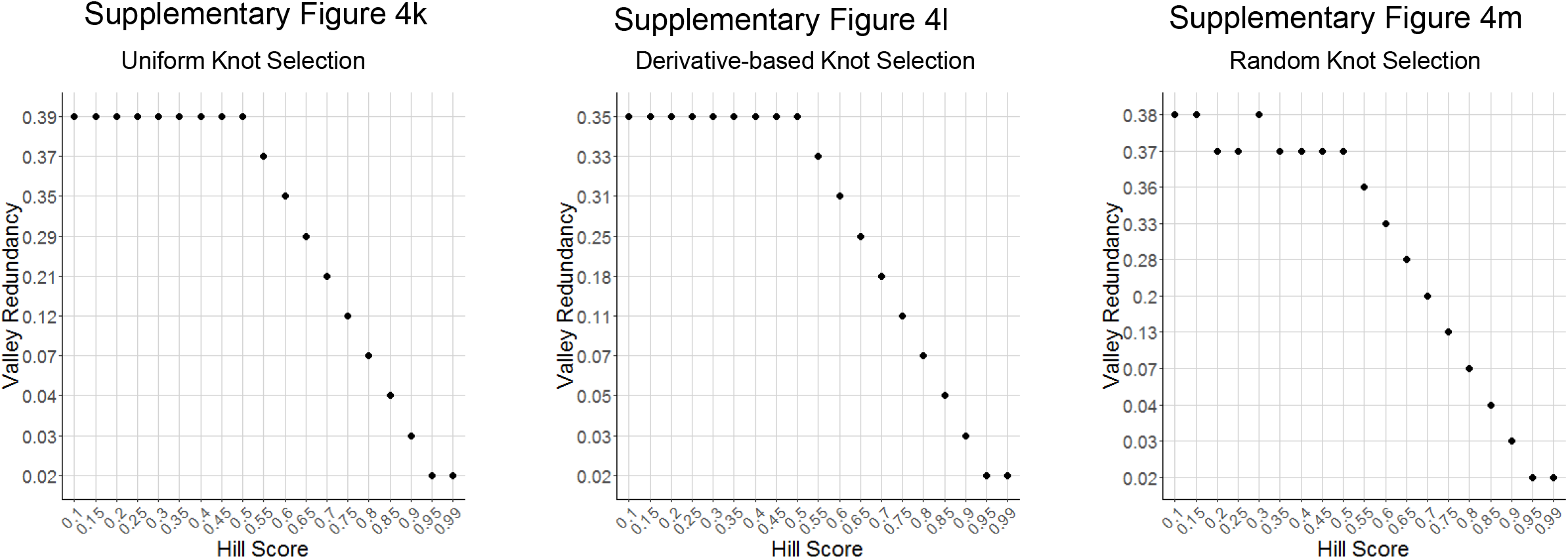

**Supplementary Figure 5.**
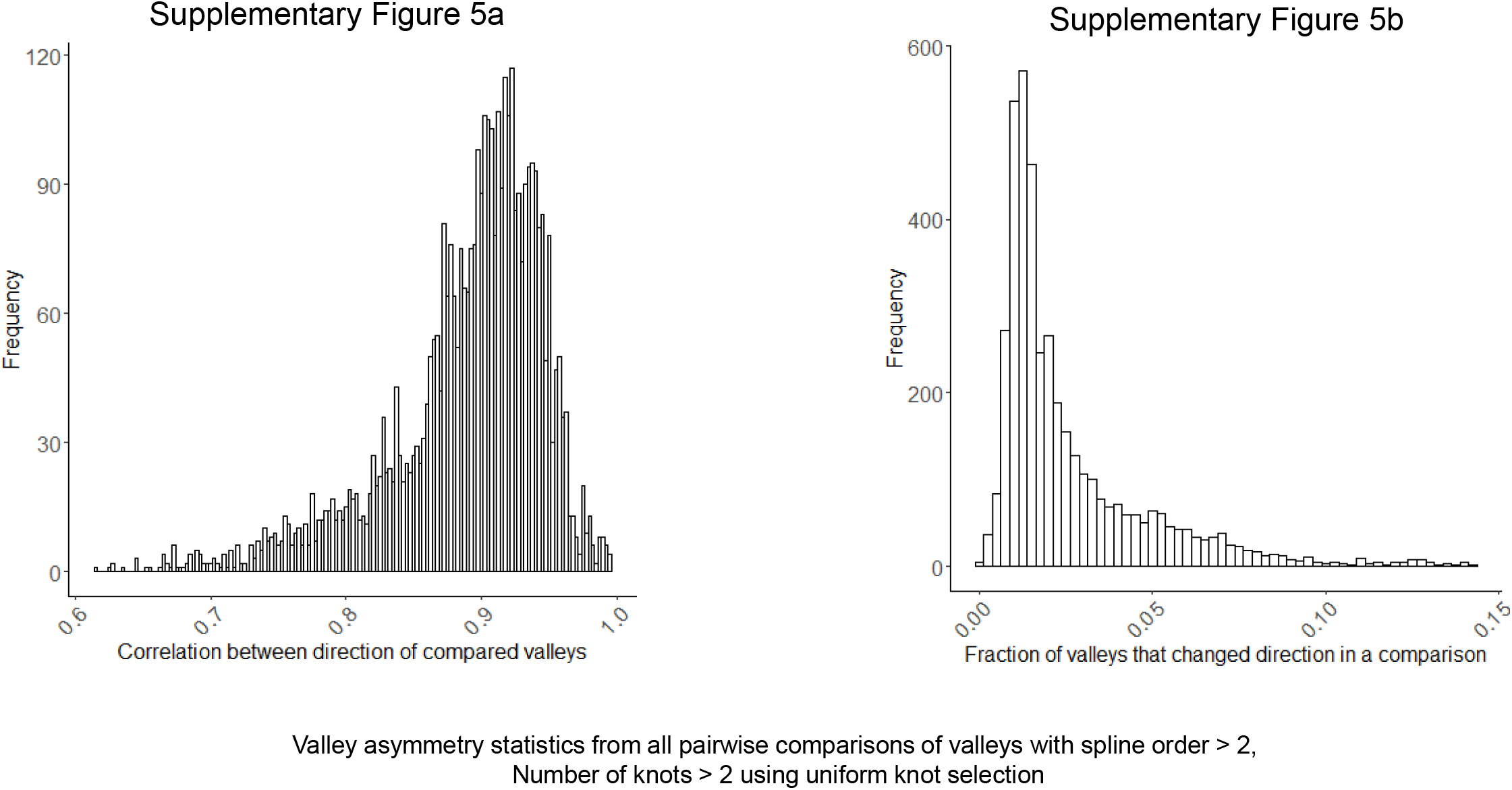

**Supplementary Figure 6.**
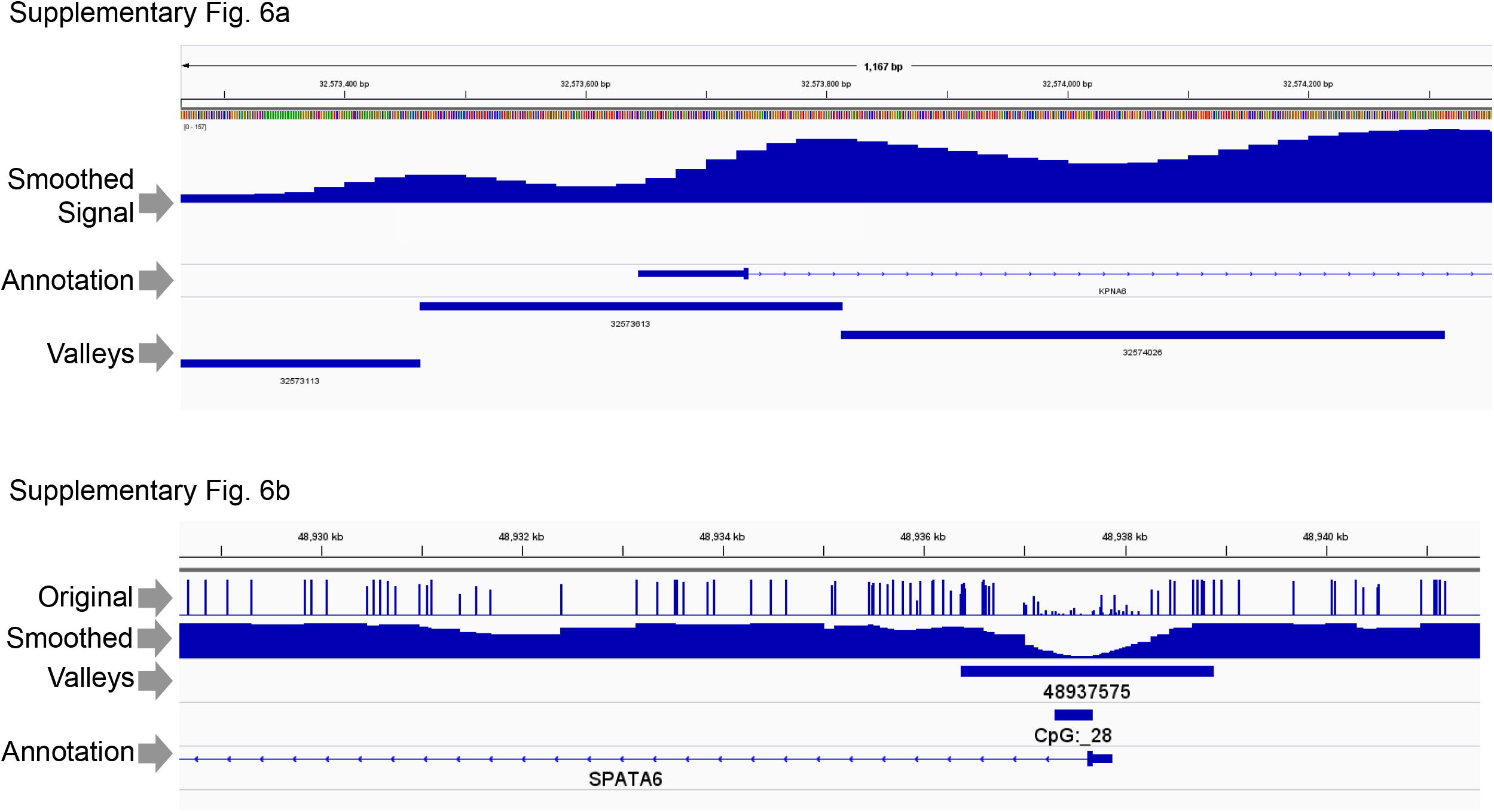

**Supplementary Figure 7.**
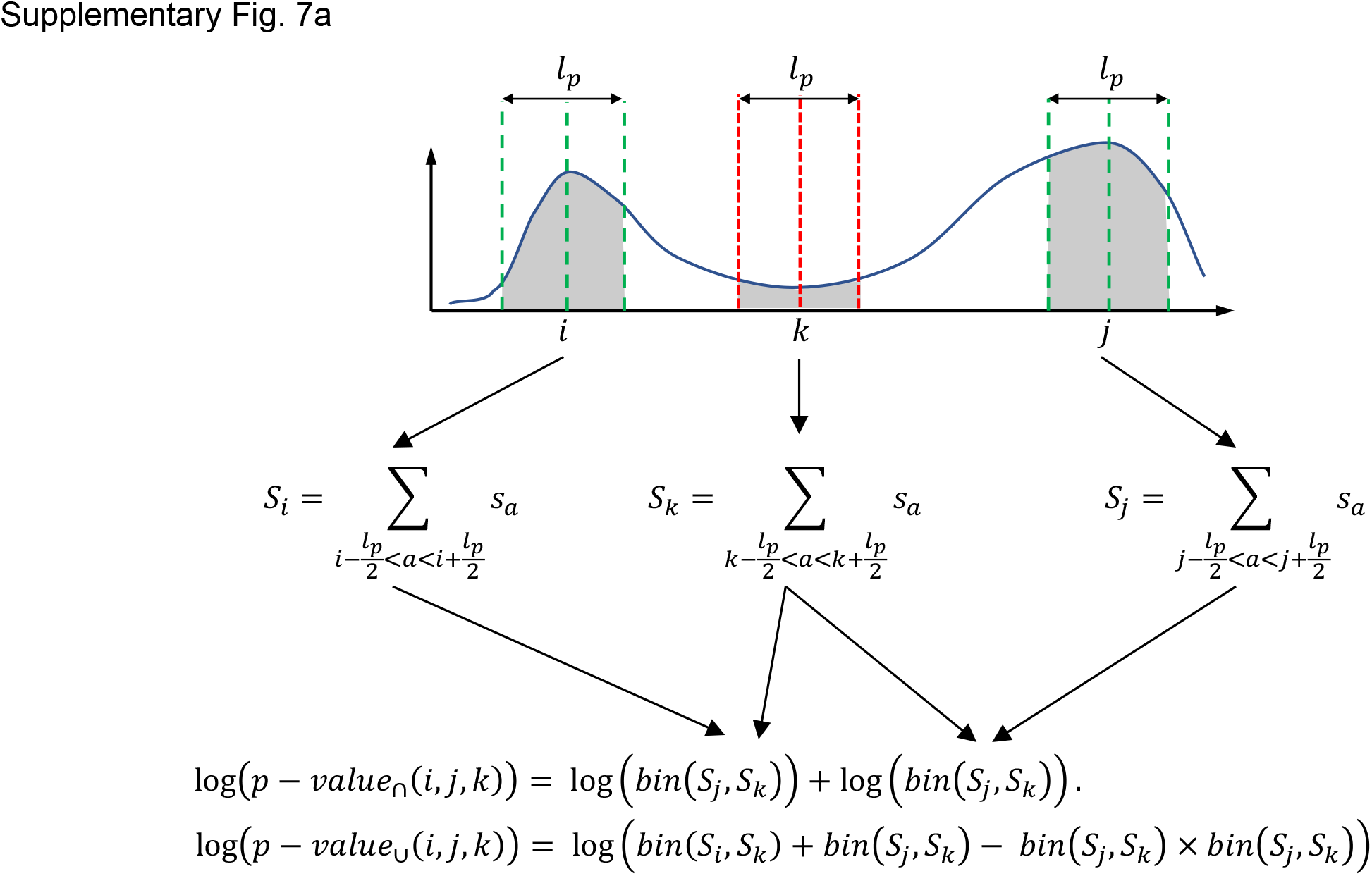
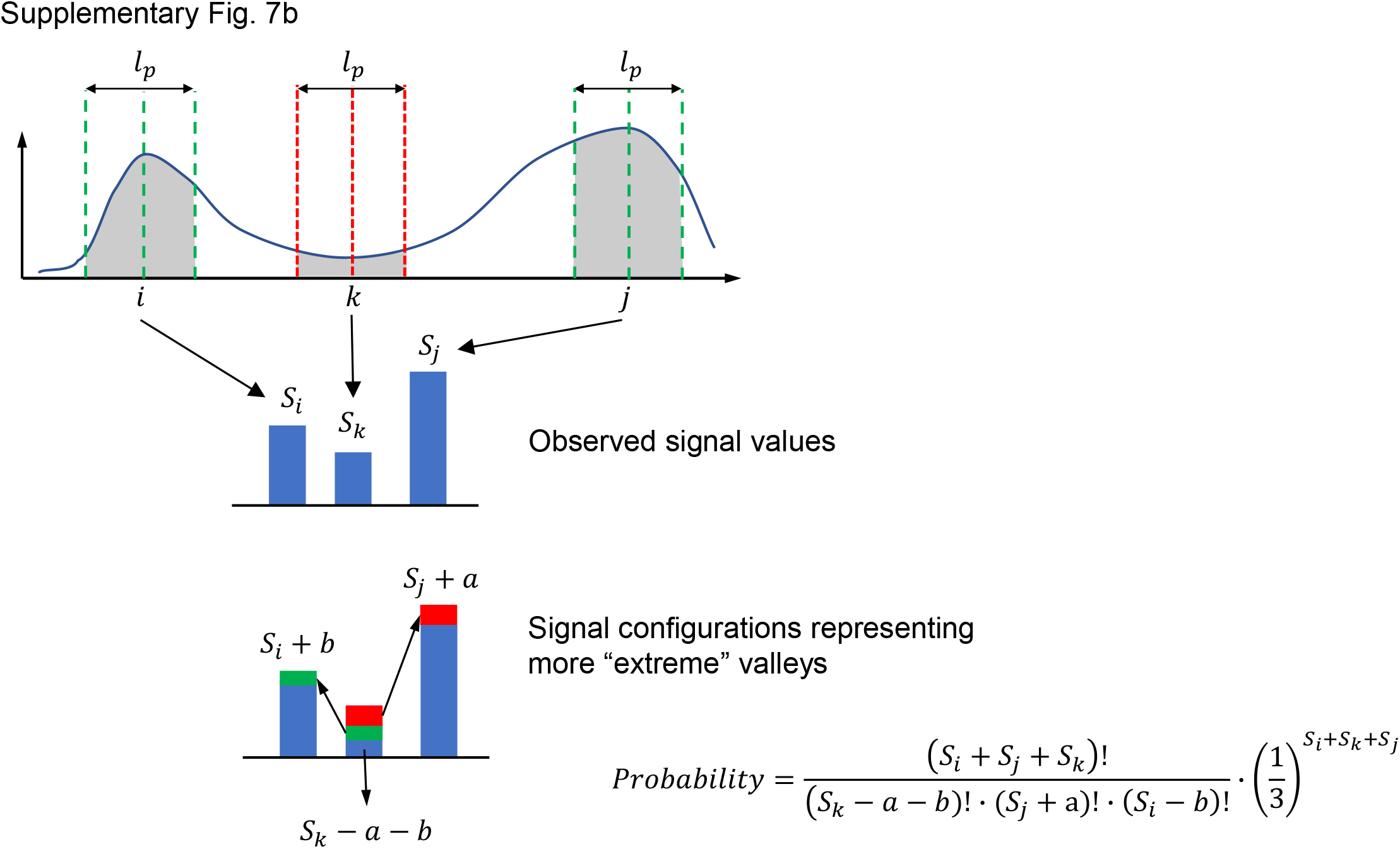
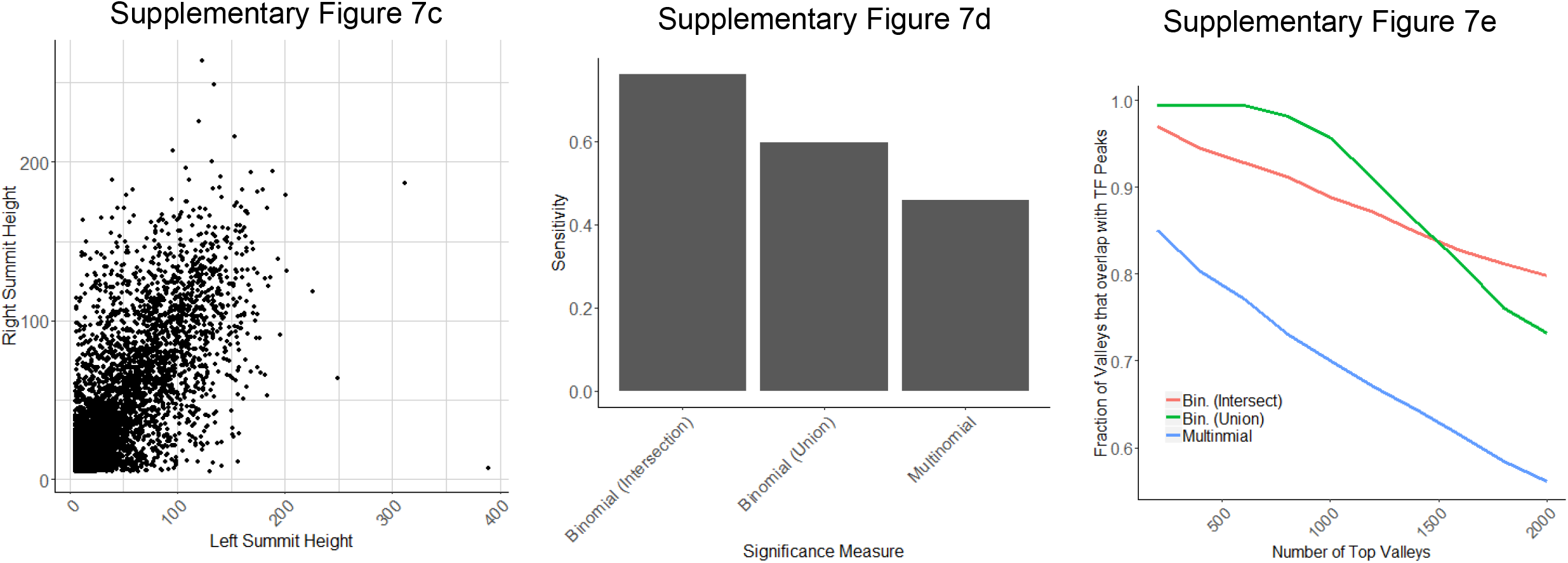

